# Predictive Ability of Enviromic Modeling in G×E Interactions for Upland Rice Site Recommendations

**DOI:** 10.1101/2025.04.05.647387

**Authors:** Marco Antônio M. Bahia, Gustavo E. Marcatti, Flávio Breseghello, Patrícia G. S. Melo, Kaio O. G. Dias, Yunbi Xu, Rafael T. Resende

**Author notes:** Corresponding author: MAMB.

## Abstract

Enviromics is an omics approach that investigates a phenomenon using all available environmental information. This study explores the use of enviromic covariates in studies of genotype × environment (G×E) interactions in upland rice in Brazil, utilizing a field trial dataset from 143 locations over 27 years, covering diverse environmental conditions. The platforms WorldClim, NASA POWER, and SoilGrids were used to extract data, resulting in 383 environmental covariates. The objective of this study was to evaluate the use of enviromic kernels to integrate GIS and genetic data for predicting upland rice productivity across Brazil and to determine the optimal number of environmental covariates required to ensure model accuracy and stability. The predictive abilities of the enviromic model peaked with around 81 covariates, stabilizing when all 383 were included, suggesting the importance of a comprehensive dataset for accurate predictions. Analysis reveals that environmental dissimilarities are more critical than geographical distance for genotypic variability, reinforcing the need to consider multiple covariates in predictive models. Heritability mapping revealed spatial variations, with regions of high heritability concentrated in southern Brazil, where genetic selection may be more efficient. The clustering of mega-environments was not efficient, highlighting the complexity of G×E interactions, and confirming that pixel-by-pixel enviromic models are a safer approach for recommending breeding actions for upland rice. This study suggests strategies to improve genotype selection for specific conditions, guiding the expansion of rice cultivation into new agricultural areas in Brazil. The findings also contribute to rice-growing regions worldwide, especially in countries cultivating upland rice under diverse conditions.

**Structured Abstract:** *Objective:* This study aimed to evaluate the predictive ability of enviromic models for site-specific recommendations in upland rice, focusing on genotype × environment (G×E) interactions by integrating environmental and phenotypic data.

*Methods:* A total of 734 field trials conducted between 1995 and 2022 across 143 Brazilian locations were analyzed. Environmental data (383 covariates) were retrieved from WorldClim, NASA POWER, and SoilGrids using GIS-based procedures. Statistical analyses included mixed linear models and Random Forest to correct for design effects, estimate heritability, and perform spatial predictions.

*Results:* Environmental dissimilarity better explained genotypic ranking than geographic distance. Predictive ability plateaued after 81 covariates, but adding more covariates reduced variance and increased model stability, supporting the use of comprehensive environmental data.

*Conclusions:* The study reinforces the need for detailed environmental characterization and the use of comprehensive enviromic models. Pixel-based predictions are more reliable than broad clustering approaches, supporting the use of virtual trials to optimize breeding strategies and resource allocation.

## 1. Introduction

Brazil is a major player in both the domestic and global markets when it comes to rice production. About 86% of the rice produced is consumed domestically, yet the country remains among the top ten rice exporters in the world (Contini et al., 2022; de Carvalho et al., 2020). The study of the interaction between rice genotypes and target environments, by incorporating the concepts of enviromics which can be defined as the study of environmental conditions that affect the development of cultivars (Resende et al., 2021), has proven promising in plant breeding (Costa-Neto et al. 2023a). The analysis of multiple environmental covariates allows the measurement of the impact of this interaction, providing insight into how rice genotypes related to the environment influence phenotypic performance (Crossa et al., 2021; Lima et al., 2017; McOmish et al., 2014). Such approaches have been applied to increase genetic gains and collect information on growth, maturation, and yield performance of rice and other crops, both in controlled environments and in the field (Resende et al. 2021a; Costa-Neto et al. 2023b).

The expansion of the upland rice crop in Brazil is of significant interest to agribusiness, considering the high demand in the domestic market and the possibility of grain export (Contini et al., 2022). In the context of developing new rice varieties adapted to the country’s climatic and soil conditions, efforts are made to optimize the genotype-environment pairing (Heinemann et al., 2024; Kanfany et al., 2021). Genotype-by-environment interaction (G×E) models, such as AMMI, GGE biplot, and factorial analysis, have been used in studies involving rice breeding (Devi et al., 2020; Hashim et al., 2021). Over the past two decades, studies involving the integration of geoprocessing with plant breeding have shown potential for rice production in non-traditional regions (Heinemann et al., 2008; Januario et al., 2018).

The technological advancement of geoprocessing in plant breeding allows for more robust environmental data collection and analysis compared to traditional climatic zoning, implemented in the mid-2000s (Annicchiarico et al., 2006; da Silva et al., 1995; Marcatti et al., 2017; Silva and Assad, 2001). Environmental data are essential for decision-making regarding the selection of suitable genotype-environment pairings, assisting in the selection of varieties adapted to specific regions (Cooper et al., 2023). The use of geoprocessing techniques offers a more precise and efficient approach to rice breeding, and its combination with enviromics techniques increases the effectiveness of selection, both for qualitative and quantitative traits, such as tolerance to adverse weather conditions and maximization of yield potential (Heinemann et al., 2021; Marcatti et al., 2017; Resende et al., 2021).

The use of enviromics aims to map information about the environment and correlate it with genotype and phenotype data, using both raw covariates and generating multiple enviromic markers (Costa-Neto et al., 2021b ; Resende et al., 2021). For the collection of environmental information, satellite images, weather station data, climate information on temperature, precipitation, wind direction and speed, air humidity, and soil physicochemical characterization, among others, can be used (Resende et al., 2024a; Xu, 2016). By combining environmental information, it is possible to integrate environmental covariates with phenotypic data, and these with genetic information from the genotypes under evaluation, facilitating the choice of the best genotype-environment combinations and predicting future scenarios (Crossa et al., 2021; Hafeez et al., 2023; Wang et al., 2024). Various authors studied the relationship between environmental covariates and genotypes to predict local yield and potential selection gains in untested environments (Resende et al., 2024b; Xu et al., 2022; Zhou et al., 2023).

Nevertheless, there is still a gap in understanding the complex interaction between rice genotypes and diverse environments, especially in non-traditional cultivation systems, such as upland rice. The lack of accurate prediction models for these areas limits the potential for including rice in sustainable systems in non-traditional areas. In this context, this study addresses this gap by integrating environmental and genetic data to enhance the selection of genotypes tailored to specific conditions in Brazil. We present different strategies for obtaining, testing, and validating environmental covariates for upland rice across the entire Brazilian territory. We will also present potential values for productivity, heritability, and associated predictive ability for the entire study area, based on the use of environmental covariates.

## 2. Material and Methods

### 2.1 Phenotypic Data

In this study, we used upland rice data from the ERBD – Embrapa Rice Breeding Dataset (Breseghello et al., 2021). These authors provide detailed information on the experimental setup and the collection of phenotypic traits. For this study, data related to genotype, location, planting date, and grain yield (GY) measured in kg.ha^−1^, were used.

A subset of 734 experiments was used, representing the period from 1995 to 2022 (27 years of the breeding program). The experiments encompass 143 distinct locations across 16 Brazilian states and the Federal District (Figure 1). Over the study period, 349 genotypes (upland rice lines and cultivars) were tested. The experiments were conducted in a randomized complete block design (RCBD), with four replications per experiment, in the context of the Value for Cultivation and Use (VCU) phase. ERBD does not include the exact geographical coordinates of the experimental locations, thus, for this study, the municipality centroid was used, as obtained from open-access repositories of the Brazilian Institute of Geography and Statistics – IBGE (2010).

**Figure 1.**
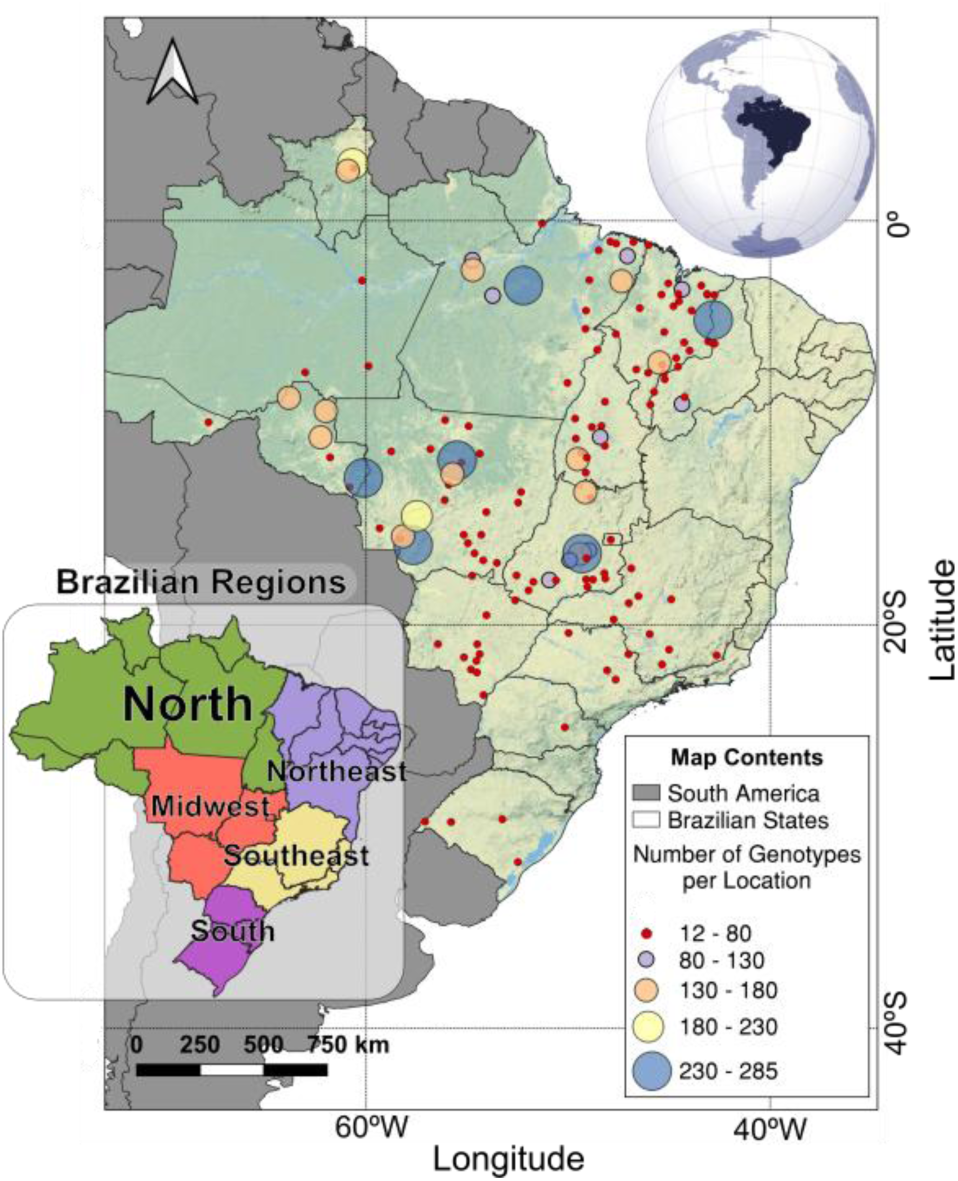
Maps of the Brazilian territory showing the geographical distribution of field trials selected for this study and the number of genotypes evaluated in Value of Cultivation and Use (VCU) trials per location over 27 years (1995-2022).

### 2.2 GIS Treatment and Acquisition of Environmental Covariates *via* Envirotyping

We configured the Geographic Information Systems (GIS) for enviromics purposes, following the steps proposed by Resende et al. (2024a). The analysis integrated grain yield data and environmental information to construct an envirome representing the combination of all available environmental covariates (Resende et al., 2021). A geospatial grid representing the target population of environments (TPE) was generated using the “Regular Points” tool in QGIS, with a resolution of 11.1 km per pixel edge (123.21 km² per pixel), covering almost the entire Brazilian territory while balancing spatial precision and computational processing limitations. The ERBD database was then concatenated with geographic coordinates from IBGE, using the municipality name as the matching key (Breseghello et al., 2021; IBGE, 2010), resulting in a dataset of 143 locations with observed phenotypic data. Environmental data were extracted for both these locations and the full grid of 87,155 points, ensuring comprehensive coverage of the study region.

The envirotypic characterization was performed based on three geoclimatic data repositories: i) WorldClim, ii) NASA POWER, and iii) SoilGrids (specifically detailed in the following items). Data was acquired automatically through Python scripts, using WGS84 as the geodetic datum.

**i. WorldClim V.2.1.** Provides high-resolution global climate data, offering monthly averages for variables such as minimum, mean, and maximum temperature, precipitation, solar radiation, wind speed, and water vapor pressure, as well as 19 derived bioclimatic variables. These datasets cover the period from 1970 to 2000 and are available in multiple spatial resolutions, ranging from 30 seconds (∼1 km²) to 10 minutes (∼340 km²), with data accessible in GeoTIFF format (sourced at: https://www.worldclim.org/).
**ii. NASA POWER.** The NASA Prediction Of Worldwide Energy Resources (POWER) project offers solar and meteorological datasets to support various applications, including renewable energy and agricultural needs. The data is sourced from NASA’s research and is provided at native resolutions; for instance, primary solar data is available on a global 1° × 1° latitude/longitude grid, while meteorological data is on a 0.5° × 0.625° grid. These datasets encompass parameters such as solar irradiance, temperature, humidity, wind speed, and precipitation, and are accessible through a RESTful API, facilitating automated data retrieval (sourced at: power.larc.nasa.govpower.larc.nasa.gov).
**iii. SoilGrids 2.0.** A global gridded soil information system that utilizes state-of-the-art machine learning techniques to predict soil properties and classes. It leverages a compilation of soil profile data and environmental covariates to produce maps of soil properties, such as organic carbon content, bulk density, cation exchange capacity (CEC), pH, and soil texture fractions, at a spatial resolution of 250 meters. These predictions are made for six standard depth intervals up to 200 cm, providing valuable information for various environmental and agricultural applications (sourced at: https://soilgrids.org/).

Data processing was conducted using a combination of Python libraries. The *rioxarray* and *rasterio* were used for manipulating, projecting, and rasterizing environmental data in TIFF format (https://pypi.org/project/rioxarray/; https://pypi.org/project/rasterio/). *Xarray* was employed to organize multidimensional data into geospatial datasets (https://pypi.org/project/xarray/), while *pandas* was used to structure tabular data and save the results in CSV format (https://pypi.org/project/pandas/). The *pathlib* library facilitated the handling of file paths and directories (https://pypi.org/project/pathlib/), and *zipfile* was used to unzip WorldClim data (https://docs.python.org/3/library/zipfile.html).

This set of tools enabled the integration and interpolation of rasterized values for each grid point using the “nearest neighbor” method. The envirotyping process resulted in 383 environmental covariates for environmental characterization and subsequent statistical analysis. The environmental covariate dictionary is provided in the Supplementary Material (Table S1).

### 2.3 Models for Adjustment and Correction of Trial and Year Effects

#### 2.3.1 General Linear Mixed Model for All Data

We fitted a linear mixed model (LMM) that includes the G×E interaction to analyze the phenotypic variability in grain yield among upland rice genotypes, adjusted for trial and year effects. This model separates genotypic effects from location and year effects, allowing for an accurate assessment of genetic performance under different environmental conditions (Figure 2, step 1 modeling). The model is described by the equation:

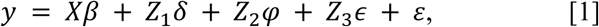

**Figure 2.**
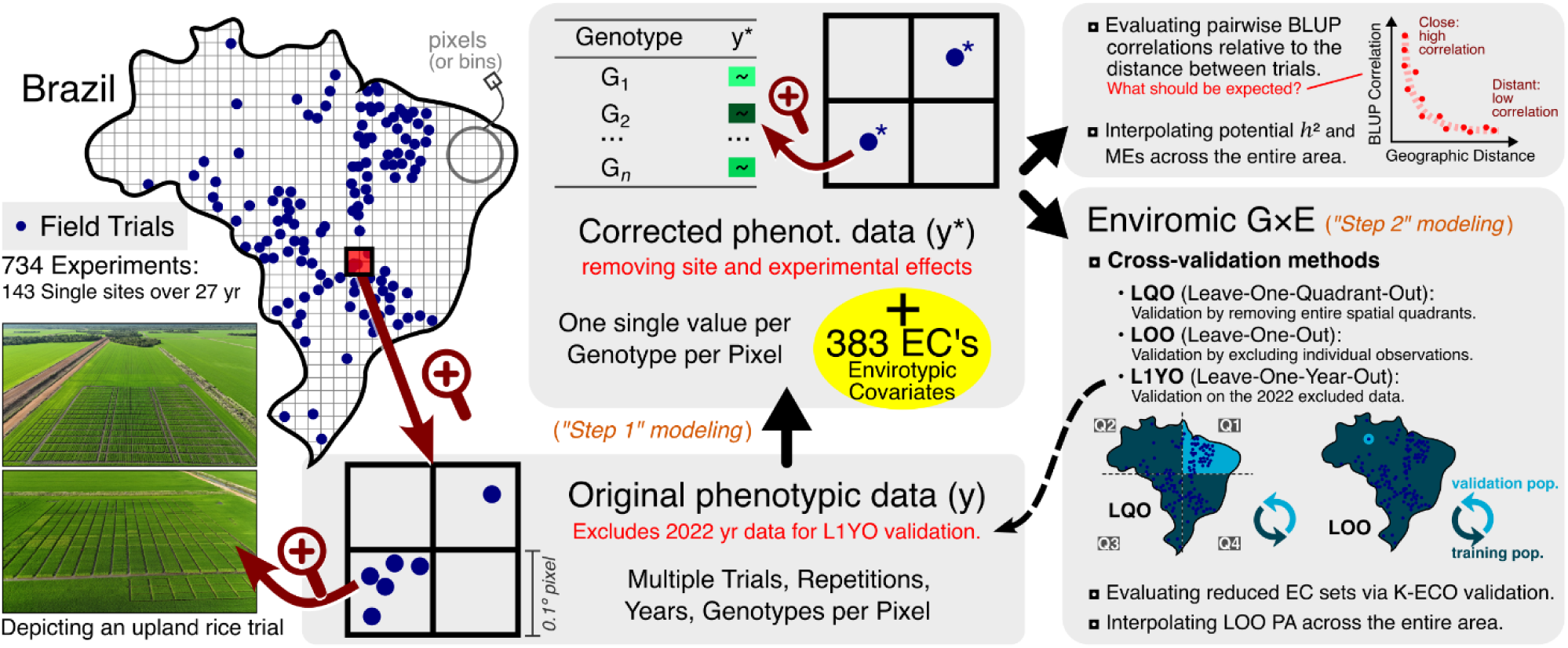
Workflow for enviromic modeling and validation in upland rice trials across Brazil. Phenotypic data is corrected for site effects and integrated with 383 environmental covariates. Four cross-validation methods (K-ECO, LQO, LOO, L1YO) assess predictive ability (PA), while interpolating estimates heritability and Mega-environments (MEs).

where 𝑦 represents the vector of observations of grain yield. 𝑋 is the incidence matrix for fixed effects, with each column corresponding to one of the 143 locations, allowing direct estimates of location effects. The vector 𝛽 contains the fixed effects associated with each location. 𝑍_1_ is the incidence matrix for the random effects of genotypes, linking genotypic effects to observed yield, while 𝛿 is the vector of random genotype effects with an identity covariance matrix multiplied by a scalar 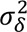. The assumption that the genotypes (𝛿) are unrelated was adopted to ensure that the model independently captures all possible genetic effects, including additive, dominance, and epistatic effects, without the influence of pre-established genealogical relationships. 𝑍_2_ relates each genotype-location combination (G×E) to the observed yields, adjusted by the ’Study Phase’, and 𝜑 captures the random effects of the genotype-location interaction, with an identity covariance matrix also multiplied by 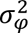. 𝑍 is used for other experimental terms such as year, trial, replication and its interactions, allowing adjustments for variations due to experimental characteristics, with 𝜖 representing the random effects associated with these terms, and an identity covariance matrix multiplied by 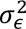. Finally, 𝜀 represents the residual errors, with an identity covariance matrix multiplied by 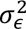, ensuring that the errors are independent and uniformly distributed.

This model was implemented using the ’lme4’ package in R software (Bates et al., 2024), which facilitates handling large datasets and effectively estimates variance components, as well as fixed and random effects. The model provides a detailed assessment of genetic variability and genotype-environment interactions while controlling for trait and year effects. The final corrected value is given by 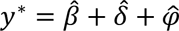, where 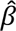 represents the fixed intercept effect—Best Linear Unbiased Estimate (BLUE)—for each location, ensuring that the response variable is adjusted to the corrected scale. This is combined with the genotypic effects—Best Linear Unbiased Prediction (BLUP)—for each cultivar (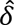); and the genotype-environment interaction effects (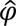) across the 143 environments where they were tested. The model described in Equation [1] also served as a baseline for predicting the genotypic values of varieties in untested locations.

#### 2.3.2 Pixel-Based Mixed Models for Local Heritability Estimation

To estimate total genetic variance and local broad-sense heritability (ℎ^2^), separate mixed models were fitted for each pixel, with independent adjustments to obtain variance components specific to each location. The general model framework previously described was used in both cases, but its formulation varied depending on the number of trials within the pixel. When a pixel contained multiple trials, the full model was applied, maintaining all terms, including genotype-by-trial interaction, to account for within-pixel variability. Conversely, when a pixel contained only a single trial, this interaction term was removed, resulting in a reduced model with different incidence matrices and dimensions, considering only genotypic and experimental effects. Despite these structural differences, both models allowed for consistent variance component estimation across spatial scales while avoiding excessive redundancy in model notation. Local broad-sense heritability (ℎ^2^) was computed as:

- When there was more than one trial in the pixel: 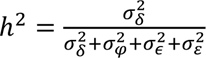
- When there was more than one trial in the pixel: 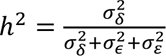

where 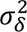 represents broad-sense genotypic variance, 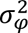 accounts for genotype-by-trial interaction variance, 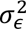 accounts for the combination of year, trial, replication and its interactions, and 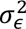 is the residual variance. The models were fitted iteratively for each pixel, and the estimated heritability values were georeferenced using the latitude and longitude of each location. These estimates allowed for spatial visualization of genetic variance across the study area, supporting the identification of regions with greater experimental precision and potential for breeding selection.

### 2.4 Genotypic Correlations across Geographical Distances and Environmental Dissimilarities

To measure the extent of genotypic correlations across different geographical distances and environmental dissimilarities, we obtained Spearman’s rank correlations of genotypic BLUE+BLUPs (𝑦^∗^) between pairs of trials (not pairs of pixels), based on the predictions generated by the model [1]. In addition, we calculated the distances between trial pairs, creating matrices for geographical distances (𝐷_𝐺_ in km) and environmental dissimilarities, which were calculated as 𝐷_𝐸_ = 𝑚𝑎𝑥(𝐸) − 𝐸, where 𝑚𝑎𝑥(𝐸) is the highest value of 𝐸, analogous to 1 minus the similarity value, assuming that the maximum similarity value will be unity. On item 2.7 of this document, we will see that ’𝐸’ is the enviromic similarity matrix and how it is obtained. This proposed approach is mathematically analogous to calculating the decay of linkage disequilibrium (LD) across the genome (Remington et al., 2001).

To model the decay curve, we used simple linear regression and loess regression (Jacoby, 2000), both implemented in the base R package. To account for semi-variance effects, weighted regressions were applied, considering the number of genotypes shared between trial pairs. Additionally, to address spatial autocorrelation, we followed the procedures described by Diniz-Filho et al. (2009). The relationship between genotypic correlations and both geographic distance (𝐷_𝐺_) and environmental dissimilarity (𝐷_𝐸_) was assessed using Mantel correlation analysis (Mantel, 1967), which quantifies the association between distance matrices and the observed genotypic correlations.

### 2.5. Predictive Modeling of Yield Using Spatial Interpolation and Machine Learning

To analyze the predictability of yield, an enclosing polygon was generated using the convex hull method with a 250 km buffer. This process ensures that predictive values are not underestimated when attempting to predict areas far from testing centers, this strategy aligns with the delimitation of target population of environments (TPEs) to enviromics context, as outlined in Cruz et al. (2025). There is a practical component to using the enclosing polygon, which is simply the suitability of the locations for the crop being analyzed.

Based on the 143 tested locations, we applied a RandomForest (Liaw and Wiener, 2002) model to predict the yield means of the pixels within the enclosing polygon. The pixel-level means correspond to the intercept (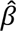) of Model 1 when fitted per pixel content, representing the GY’ BLUE of the pixel. The spatial interpolation technique of ordinary kriging was used, along with the definition of a theoretical variogram through the spherical model. To verify the predictive ability, a correlation was made between the observed data in the 143 locations and the data predicted by the RandomForest model (Li et al., 2022; Medeiros et al., 2024).

### 2.6. Interpolating Mega-Environments

Two steps were necessary for the generation of mega-environments. First, hierarchical Ward clustering was performed, similar to the method used by Rajak et al. (2024), aiming to minimize the sum of squared costs within clusters, also minimizing the internal variance of each cluster. The clustering was performed using the “hclust” function from R’s “stat” package, with the ’ward.D2’ option (Bolar, 2022). The cost function employed is expressed as 1 minus the correlation between the experimental sites taken in pairs. The ideal number of groups was visually determined using the ’Elbow Curve’ method (Shi et al., 2021), with the criterion of creating relatively balanced groups in terms of the number of test sites. The most satisfactory results were obtained for 5 and 9 clusters, and the grouping into 5 clusters were selected for subsequent analyses.

Next, we performed the interpolation of clusters across the Brazilian territory using a weighted k-nearest neighbor classifier with a ’Gaussian kernel’ function, employing a specific interpolator for categorical variables. This process was executed using the kknn function from the kknn package in R (Schliep et al., 2016).

### 2.7. Enviromic Model *via* Kernels

The kernel is a mathematical function that computes the similarity between two data points – in this case, environments - based on the available environmental covariates (Costa-Neto et al., 2021b). It allows the transformation of the input data into a higher-dimensional space, capturing complex relationships between environments that may not be apparent in the original space. In this study, the kernel matrix 𝐸 is constructed from the matrix of envirotypic covariates 𝑀. Specifically here, raw environmental covariates were used as enviromic markers, totaling 383 markers. The kernel matrix 𝐸 was calculated as the product of 𝑀 by the transpose of 𝑀 (𝑀 × 𝑀^𝑇^), normalized by the trace (𝑡𝑟()) of 𝑀 × 𝑀^𝑇^, divided by the number of rows in 𝑀 (𝑛). This normalization ensures that the kernel matrix 𝐸 efficiently captures the similarities between environments based on enviromic markers. The equation to calculate 𝐸 is (Jarquín et al., 2014):

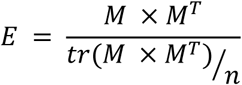

To predict genotype by environment interactions, the structure is defined, according to Jarquín et al. (2014) and Costa-Neto et al. (2021a), as follows (Figure 2, step 2):

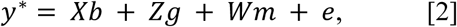

where: 𝑦^∗^ represents the vector of corrected phenotypes, as described above. 𝑋𝑏 describes the fixed effects, with 𝑋 being the design matrix for fixed effects, and 𝑏 the vector of coefficients for the trial effects. 𝑍𝑔 represents the random genetic effects, with 𝑍 being the incidence matrix linking individuals to their random genetic effects 𝑔, with distribution 𝑁(0, 𝐺 × 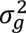), where 𝐺 is the identity matrix and 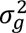 is the genetic variance. 𝑊𝑚 includes the random effects of enviromic markers, with 𝑊 being the incidence matrix and m the associated random effects, with distribution 𝑁(0, (𝐸 ⊗ 𝐺) 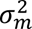). 𝐸 is described above, and 𝑒 is the vector of residual errors, with distribution 𝑁(0, 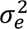). This model not only facilitates the estimation of main genetic and environmental effects but also allows for a detailed analysis of phenotypic variability under different environmental conditions.

#### 2.7.1. Enviromic Predictive Ability of Genotypic Values for Untested Locations

The first cross-validation strategy aimed to evaluate the impact of the number of ECs on predictive enviromic abilities, with a particular focus on K-fold Environmental Covariates Out (K-ECO). In this approach, a k-fold cross-validation scheme (k = 10) was applied, where 90% of the trials were used for training and 10% for validation, while simultaneously varying the number of ECs included in the model. A total of 10,000 different EC combinations were tested, ranging from reduced subsets to the full set of 383 ECs, ensuring that multiple validation groups were explored even when all ECs were included.

Additionally, three other cross-validation strategies were implemented: (i) Leave-One-Quadrant-Out (LQO), where models were validated across virtual quadrants, defined solely based on the median latitude and longitude of the experimental data, without considering bioclimatic or environmental criteria (Figure 2); (ii) Leave-One-Out (LOO), in which a single location was left out and predicted; and (iii) Leave-One-Year-Out (L1YO), where models trained on data from 1995 to 2021 were used to predict 2022, which remained completely excluded from the modeling process (Rogers and Holland, 2022).

Throughout all validation process, Pearson correlations were computed for each location to prevent confounding effects, such as those that could arise from Simpson’s paradox (for more details, see Pearl (2014)).

To understand where the Enviromic model has the potential to perform better or worse across the study area, we applied a RandomForest model to interpolate the LOO validation results. This spatial interpolation provided a continuous estimate of predictive ability, highlighting regions where the model performs more reliably based on environmental patterns.

## 3. Results

The geographical distribution of the data is presented in Figure 1, where the 143 locations tested were classified according to the number of upland rice genotypes evaluated. Approximately 4.2% of these locations were tested each with more than 200 genotypes. In contrast, in major irrigated rice-producing centers located in the Southern Region of Brazil, the number of upland rice genotypes evaluated per location was significantly lower, not exceeding 21 genotypes at any location.

The Southeast Region has only 13 experimental locations, with the number of genotypes analyzed not exceeding 50 per location. In the Northeast, 31 test locations were recorded, with the number of genotypes evaluated ranging from 16 to 233 and a median of 28 genotypes per location. In the Northern Region, 41 locations were used for experimentation, with the number of genotypes analyzed ranging from 16 to 271, presenting a median of 60 genotypes, with a concentration in areas of agricultural expansion. The Central-West Region concentrated 53 test locations, representing 37% of the total, had the number of genotypes tested varying widely between 12 and 285, and a median of 40 genotypes per location.

To validate genotypic predictions, the Brazilian territory was divided into four virtual quadrants based on the median of latitude and longitude data (Figure 2). Quadrant 1 (Q1) included 51 testing locations (35.66%), while Quadrant 2 (Q2) covered only 19 (13.2%). Quadrants 3 (Q3) and 4 (Q4) grouped 41 and 32 testing locations (28.67% and 22.37%), respectively. The number of genotypes tested per quadrant was relatively balanced, with 314, 291, 325 and 316 for Q1 to Q4, respectively.

The mixed model was fitted using data from all years and locations, assuming genotype, environment, and genotype-by-environment interaction as random effects. The residual variance was 475,363 (54.2% of the total), followed by the environmental effect with 1,049,522 (11.9%), the genotype-by-environment interaction with 159,694 (18.2%), and the genotypic effect with 63,943 (7.3%). These results are illustrated in Figure 3, which presents the predicted grain yield for each trial in the training period (1995–2021, Figure 3-A) and in the validation year (2022, Figure 3-B). To facilitate interpretation, five genotypes were highlighted: BRS Primavera, the first long and slender grain cultivar developed by Embrapa; BRS Bonança, BRS Sertaneja, and BRS A502, also from Embrapa, are recognized for their grain quality and lodging resistance; while Cambara is a well-recognized cultivar from a private company.

**Figure 3.**
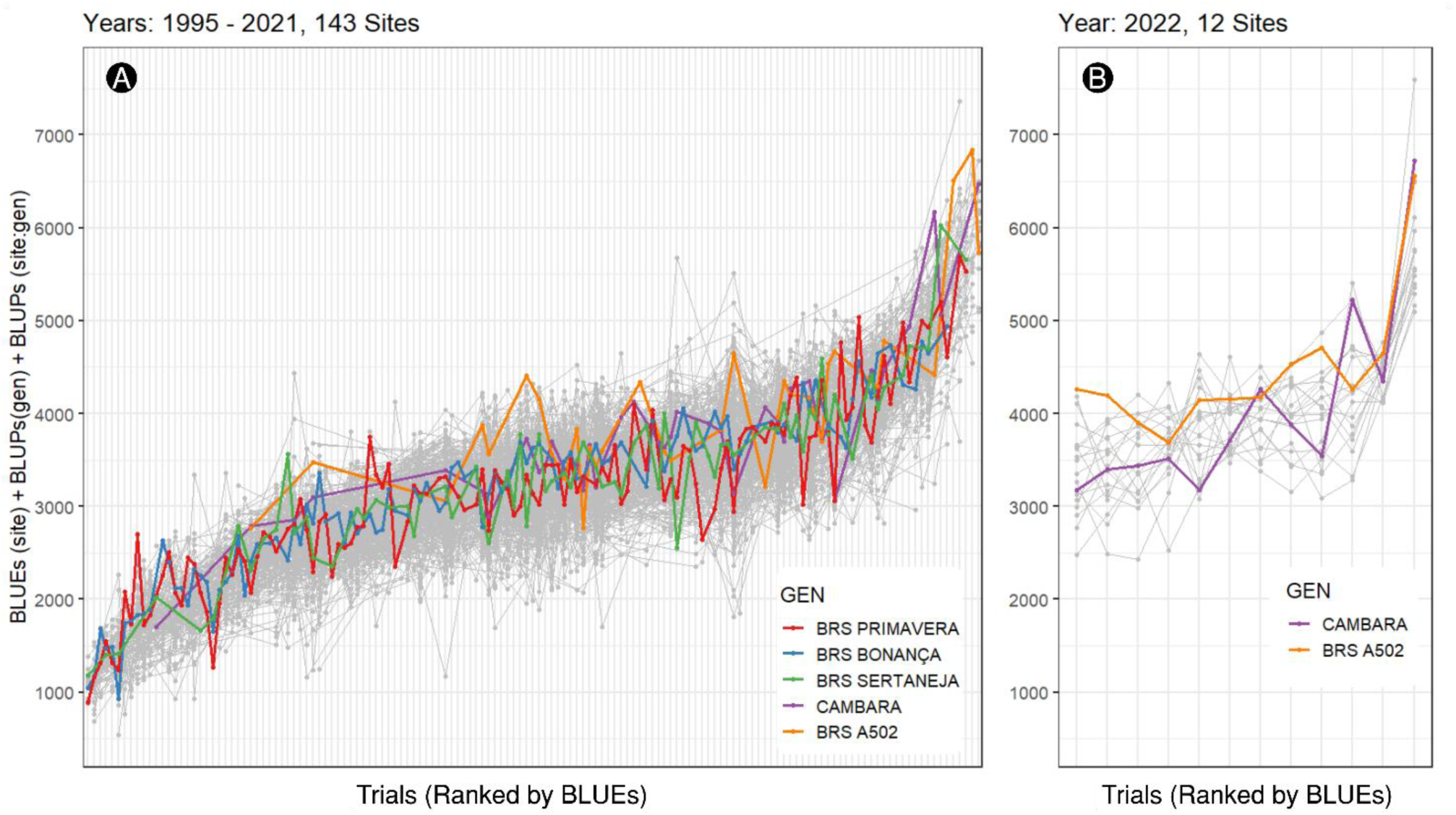
Grain yield prediction by trial for training years (A) and for the valid year (B). Colored lines emphasize key genotypes.

The modeling strategy adopted in this study (Equation 1) is based on a unified model structure, which aggregates several experimental design components into a single random term. This approach simplifies model specification while retaining the key sources of variation. The similarity between the used model and the full one is illustrated in Supplementary Material Figure S1, where predictions, residuals, and variance components show high correspondence between both models. Supplementary Material Table S2 presents the variance components from the full model, where the interection between the experimental location and the time was the most expressive source (50.9%). Although the interaction between genotypes and time was statistically significant (p < 0.01), it accounted for only 0.68% of the total variance. This limited contribution is also visually confirmed in Supplementary Material Figure S2, where genotypic performance across years shows minimal re-ranking.

We used a pairwise matching approach to assess the genotypic correlation across experimental locations, calculating the correlation of genotypic rankings between pairs. This correlation was then associated with the geographical distance and the environmental dissimilarity between locations. The average distance between the trials was 1,215 km, ranging from 5.5 km to 3,924 km. Using the Mantel test to correlate geographical distances with genotypic rankings (Figure 4-A), no significant values were found (p-value: 0.6535). This result suggests that geographical distance alone does not have a direct impact on similarity of genotype performance between locations.

**Figure 4.**
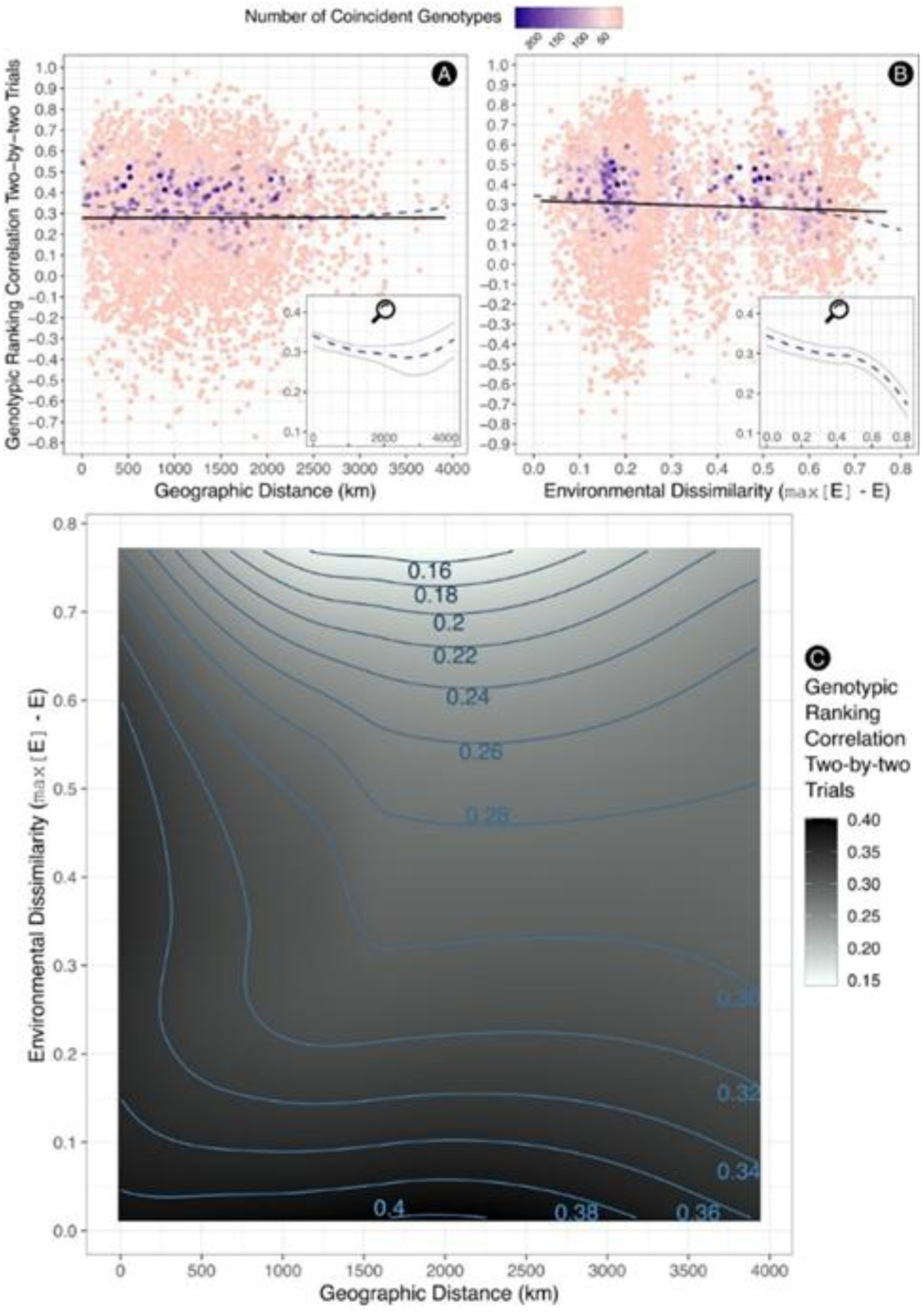
Scatter plot between two matrices: **A)** Genotypic correlation matrix × distance matrix; **B)** Genotypic correlation matrix × environmental dissimilarity matrix constructed from Kernel matrices using environmental covariates; **C)** Environmental dissimilarity × geographical distance × genotypic ranking correlation. In parts A and B, the intervals between 0.2 and 0.6 of the genotypic correlation are highlighted for better visualization of the curve behavior. Each point represents a combination between two tested locations, and the color gradient (from pink to purple) indicates how many coincident genotypes are present in that combination.

The correlation between environmental dissimilarities, derived from the environmental covariates, with genotypic ranking correlations between pairs of locations (Figure 4-B), was significant (p-value: 0.0099). This indicates that environmental dissimilarities played a more relevant role than geographical distance in the observed genotype performance between locations. Furthermore, by combining the analyses of geographical distance and environmental dissimilarity (Figure 4-C), a clear trend was observed: locations that exhibit low environmental dissimilarity and are geographically close tend to show greater equivalence in genotypic rankings. This suggests that, in addition to geographical proximity, the similarity of environmental conditions must be taken into consideration in multi-environment testing networks.

The K-ECO validation results showed that the predictive ability of the enviromic model increases with the number of ECs (Figure 5-A), reaching a stabilization point around 81 ECs, beyond which additional covariates provided minimal improvement. ECs were randomly divided into training and validation sets, in a proportion of 80% and 20%, respectively. Model accuracy peaked at 0.536 with approximately 81 covariates, followed by a plateau up to the total of 383 ECs. No single EC or small subset performed well in isolation. As shown in Figure 5-B, the amplitude of predictive ability decreases as the number of ECs increases, indicating greater model stability. Additionally, Figure 5-C demonstrates that predictive ability is associated with the model-explained G×E variance, suggesting a structured relationship between model performance and environmental variability. A saturation point is observed, beyond which additional explanation of G×E variance results in a decline in predictive ability.

**Figure 5.**
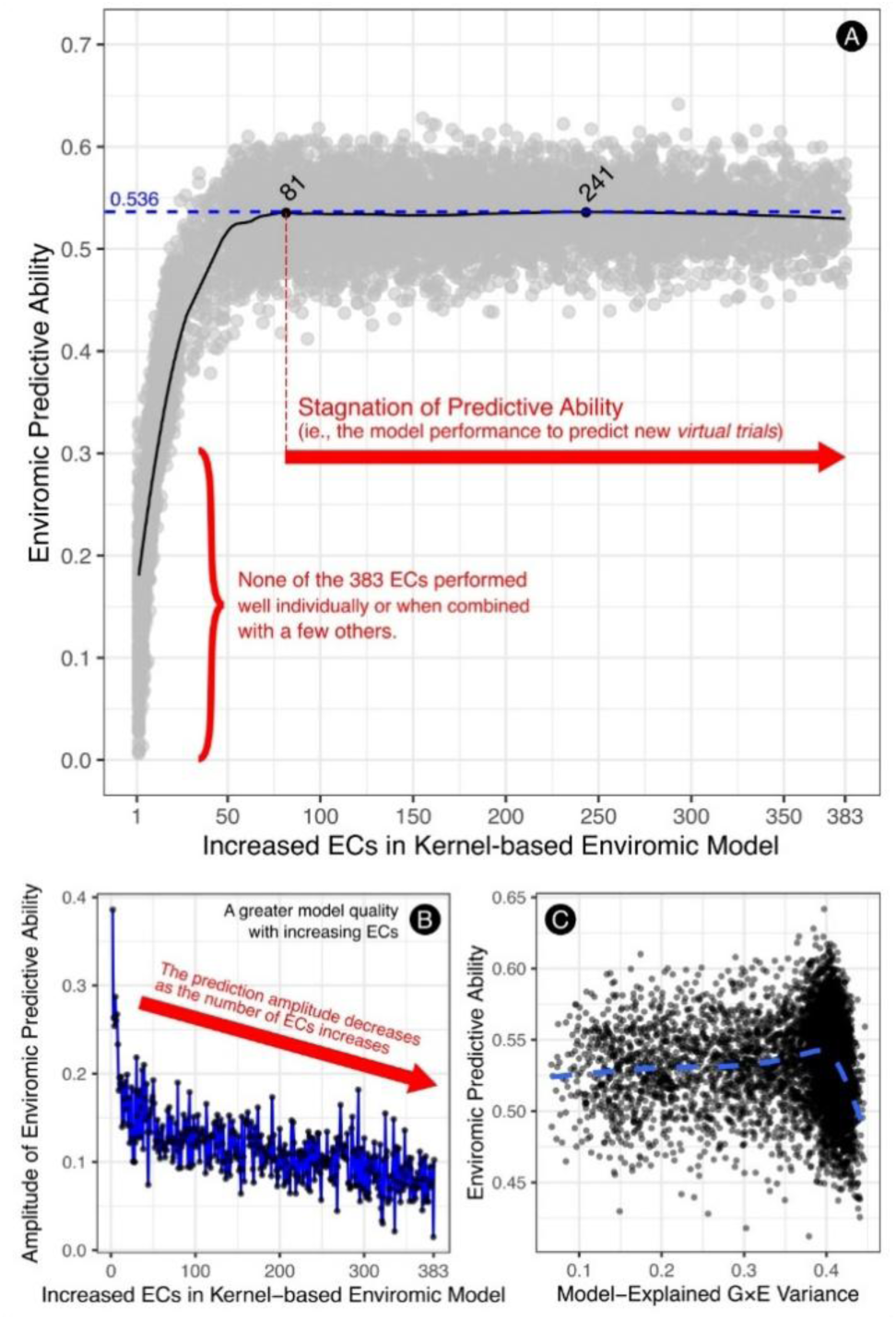
Predictive ability using environmental covariates. **A)** Effect of the number of covariates on the predictive ability of enviromic models using raw environmental covariates; **B)** Decrease in the range of predictive ability versus number of environmental covariates in the model kernel; **C)** Relationship between genotype × environment interaction explained by the model and predictive ability of the enviromic model.

A summary of the comparison between three other methods for estimating predictive abilities is shown in Table 1, contrasting a conventional Genotype × Environment interaction model and a Kernel model with environmental covariates. The first method used, validation across quadrants, showed a predictive ability of 8.2% for the traditional model and 51% for the enviromic marker-based model.

**Table 1.** Three different strategies (LQO, LOO, L1YO) for obtaining Predictive Abilities (for all genetic materials in new environments) comparing a traditional G×E model and the Kernel model using environmental covariates (ECs).

| Strategy | Valid Set | ngen | Baseline G×E method |  |  | Enviromics Kernel w/ ECs |  |  |
| --- | --- | --- | --- | --- | --- | --- | --- | --- |
|  |  |  | PA | rmse | rmse% | PA | rmse | rmse% |
| LQO | Q1 | 314 | 0.065 | 836.25 | 28.88 | 0.486 | 306.75 | 9.78 |
|  | Q2 | 325 | 0.141 | 834.78 | 35.04 | 0.511 | 306.63 | 10.88 |
|  | Q3 | 291 | 0.080 | 877.41 | 32.49 | 0.572 | 280.09 | 8.86 |
|  | Q4 | 316 | 0.035 | 1125.32 | 41.04 | 0.510 | 370.59 | 11.80 |
|  | Total | 349 | **0.082** | **903.96** | **33.76** | **0.510** | **317.01** | **10.41** |
| LOO |  | 349 | **0.064** | **939.39** | **35.35** | **0.542** | **307.14** | **10.12** |
| L1YO |  | 12 | **0.428** | **456.43** | **11.44** | **0.529** | **411.02** | **10.33** |

The LOO method presented the greatest discrepancy between results, with a predictive ability of 6.4% for the traditional method and 54.2% for the model with environmental covariates; while the L1YO validation showed the most balanced result, with 42.8% and 52.9% for the respective tests. The results obtained for the Kernel model with covariates are similar to the values found when using environmental covariates with training and validation groups in an 80%-20% ratio (Figure 5-A).

Based on data of grain yield (GY), heritability, and predictive ability from the 143 test locations, an interpolation of these characteristics was performed for an enclosing polygon within the Brazilian territory. Projected GY potential (Figure 6-A) varied within a range from 2000 to 4000 kg.(ha.year)^−1^, with the lowest values concentrated in the region bordering Paraguay, while the high values were observed in various areas of the Southeast, North, and Northeast regions. Most of the territory showed projected values above 3000 kg.(ha.year)^−1^. The correlation between observed and projected GY at 143 test locations was 0.606 (p-value < 2.2e-16).

**Figure 6.**
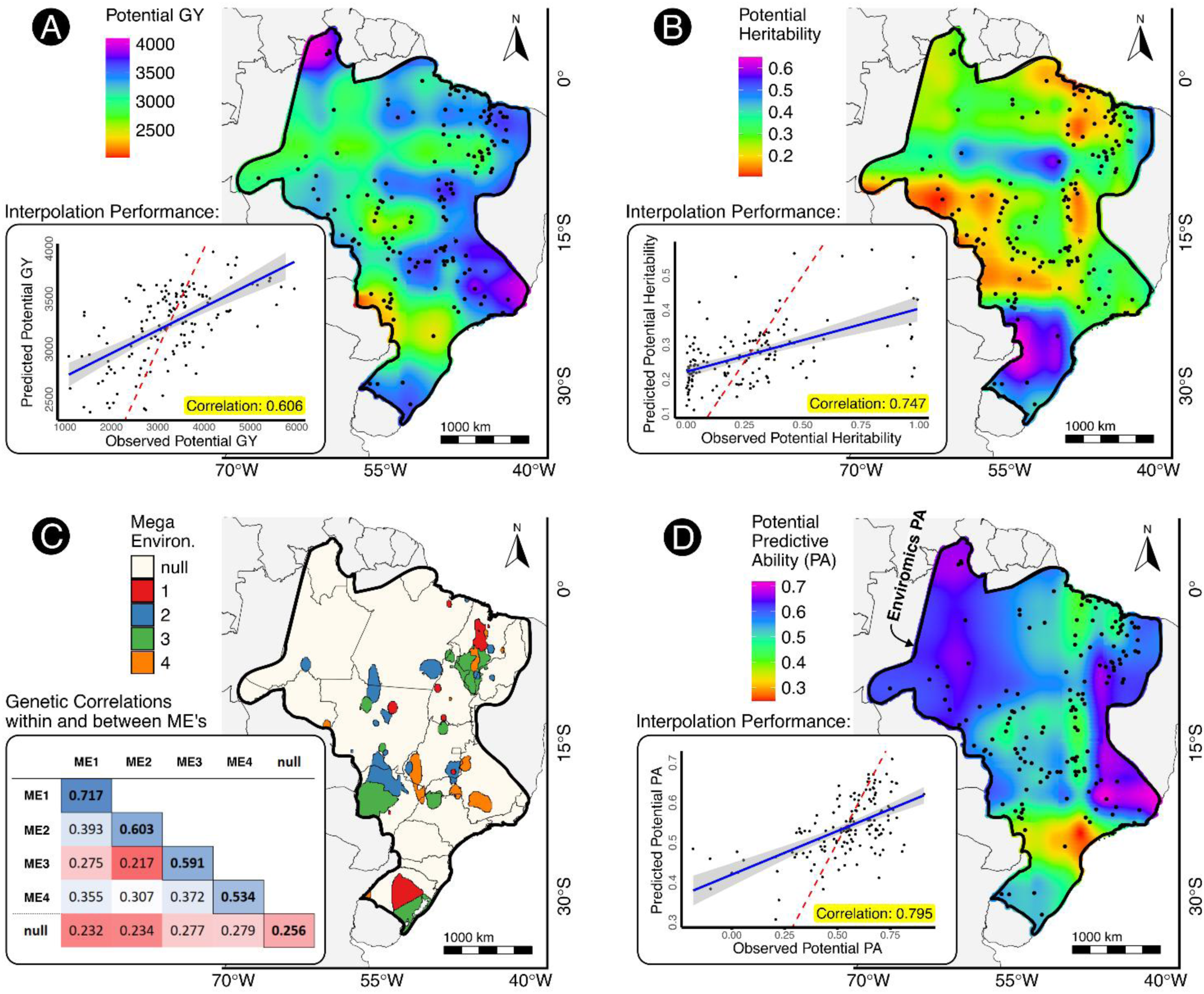
Estimated potential values within the enclosing polygon: **A)** Grain Yield and **B)** Heritability. Each map presents the linear correlation between the sample values and the values estimated by the applied model. **C)** Breeding Zones across the Brazilian territory, based on genetic correlations among test locations. **D)** Predictive Ability map showing the expected accuracy of phenotypic predictions across the region.

Potential heritability data ranged from 0.1 to 0.6 (Figure 6-B), with areas of high heritability concentrated in the South of the country. Most of the map showed heritability values between 0.25 and 0.35, with a correlation of 0.754 between the observed and projected data. Potential predictive ability (PA), shown in Figure 6-D, was higher in scattered regions of the Southeast, North, and parts of the Central-West, with values predominantly above 0.6, suggesting high reliability of the predictions in these locations.

As for the Mega-environments (Figure 6-C), the Ward hierarchical clustering applied to the 143 locations resulted in the formation of five distinct groups. Most of the territory was not grouped into any of the defined ME, forming a non-grouping zone. ME1 and 3 showed a dispersed distribution, with clusters both in the North and South of the country, while ME2 and 4 were more concentrated in specific regions. The lowest aggregation value was observed in ME4, with a minimum genetic correlation of 0.534, while Zone 1 had the highest minimum aggregation value, with a correlation of 0.717. ME3 was the most divergent among the other MEs, showing an average correlation of only 0.288 with the others.

The spatial interpolation of Potential Predictive Ability (PA) shows a heterogeneous distribution across Brazil, with higher PA values (>0.6) in the North and parts of the Central-West and lower PA values (<0.3) in the South and Southeast (Figure 6-D). The interpolation performance, evaluated by the correlation between observed and predicted PA, reached 0.795, indicating a strong relationship between the modeled and empirical data.

## 4. Discussion

### 4.1. Analysis of Genotypic Adaptation for Upland Rice in Brazil

In a multi-environment trial as the VCU, experimental locations are used to test the plasticity and stability of genotypes under relevant environmental conditions within the target population of environments (Cooper et al., 2023). Although some of these areas may not be suitable for commercial cultivation due to legal, geomorphoclimatic, or logistical reasons, they are useful for understanding theoretical responses of upland rice genotypes across environmental gradients.

The geographical distribution of the breeding trials (Figure 2) shows broad spatial coverage, highlighting the importance of considering environmental factors across spatio-temporal variations in the analysis of genotypic performance. This underscores the need for predictive models that integrate these diversities and complexities to generate more accurate and realistic predictions, as presented by Resende et al. (2021); Rogers and Holland (2022). The use of climate zoning and precision agriculture models that rely solely on geographical proximity is inadequate, as they overlook the environmental covariates that directly influence the phenotypic potential of a genotype (Resende et al., 2024b).

The mixed model fitted to the entire dataset accounted for the main components of phenotypic variance. The residual variance represented 54.2% of the total, followed by environmental effects (11.9%), genotype-by-environment interaction (18.2%), and genotypic effects (7.3%). These components are illustrated in Figure 3, which displays the predicted grain yield across trials for the training years (1995–2021) and the validation year (2022), along with the reaction norms of selected genotypes across environments. The variation in trajectory patterns across sites reflects the magnitude of G×E interaction.

For the data analyzed in this study, using Mantel correlation, no relationship was found between the geographic distance between trials and genotypic correlation, as presented in Figure 4-A. Significant results were found, however, when genotypic correlation and environmental dissimilarity between locations, calculated from environmental covariates, were analyzed (Figure 4-B). Although it is common to apply models that take into account the Euclidean distance between environments, it may be more effective to use models that incorporate environmental covariates (or enviromic markers) to determine environment similarity (Costa-Neto and Fritsche-Neto, 2021; Resende et al., 2021).

Genotype rankings are correlated between locations near to each other, provided that environmental dissimilarity between them is low (Figure 4-C). This suggests that cultivar recommendation should consider not only geographical distance or the representativeness of administration regions, as environmental similarity between locations plays an even more important role in predicting cultivar’s performance.

### 4.2. Concerns When Using Few Environmental Covariates in Enviromic Prediction

The enviromics approach can accelerate the early stages of plant breeding by using phenotypic data from consolidated trials to make better genotypic selections based on the characterization of the local envirome. The parameters used in climate zoning and precision agriculture remain valid but exhibit a low level of detail regarding the environmental vertex of the breeding triangle (Crossa et al., 2021). Using insufficient environmental covariates for phenotypic prediction can introduce inaccuracies and errors within the model, decreasing reliability in genotype-by-environment (GxE) interections.

The importance of adequately characterizing GxE interactions is highlighted by Rebollo et al. (2023), to improve the accuracy of predictive models in rice breeding programs. The study emphasizes the use of random regression models with climatic variables as a promising approach to handling G×E, especially in scenarios of prediction in untested environments. These models showed a significant increase in predictive ability in both tested and untested environments, supporting the idea that a good predictive model should be holistic and include the full range of available environmental covariates (Figure 5-A).

Similarly, the results of the present study indicate that the saturation point for the model’s predictive ability was reached at 81 environmental covariates, with predictive accuracy stabilizing at above 0.5 (Figure 5-A). Beyond this point, there were no further accuracy gains, however, the inclusion of all covariates was beneficial in stabilizing predictive ability results. Figure 5-B shows a decrease in the variability of predictions as more covariates were added to the model. This behavior reinforces the importance of considering as many environmental covariates as possible in model construction, ensuring not only more accurate predictions but also greater consistency. In this sense, enviromic prediction is similar to genomic prediction, where a large number of molecular markers are used to predict breeding values, irrespective of the significance of each marker (Chen et al., 2023; Fernandes et al., 2024).

Lau et al. (2023) emphasize the importance of methods such as bootstrap aggregating (bagging) and Random Forest for dealing with complex interactions between genetic and environmental factors, demonstrating greater statistical power in detecting GxE interactions and avoiding the loss of information that occurs when using few variables (Figure 5-A). Similarly, Li and Gutierrez (2023) reinforce that the inclusion of multiple environmental covariates in prediction models not only increases the accuracy of phenotypic predictions but also ensures that subtle interactions between the environment and genotypes are efficiently captured.

It is noteworthy that both predictive accuracy and the variance explained by genotype-environment (G×E) interactions in the model reach a peak (Figure 5-C). This peak suggests that there is an optimal point of G×E variance explanation by the model, beyond which predictive accuracy decreases. When the model explains more than 40% of the total variance in G×A interactions, predictive ability tends to decline, indicating that beyond this point, optimizing the explained variance does not necessarily translate into better predictive performance, although it remains important for maintaining consistency in predictions. This phenomenon, as indicated by Resende et al. (2024a), is also common in enviromic predictions. Caution is needed when selecting a fixed number of environmental covariates solely with the aim of maximizing the explanation of G×A interactions, as this can result in a significant decrease in predictive ability.

### 4.3. Phenotypic Interpolation in Diverse Environments and Validation Methods

The statistical analysis of the prediction data for GY and heritability can provide a foundation for improving the initial stages of upland rice breeding programs (Hasan-Ud-Daula and Sarker, 2020; Zhao et al., 2022). The application of the Random Forest model allowed the prediction of genotype performance in untested environments based on test locations, using both phenotypic data and environmental covariates. As discussed by Sekulić et al. (2020), machine learning techniques such as Random Forest have shown better results than traditional spatial interpolation methods (e.g., kriging), especially when incorporating environmental covariates. The interpolation of potential grain yield (Figure 6-A) indicated more suitable areas for upland rice cropping, particularly in the Southeast and far North of Brazil.

An interpolated heritability, in turn (Figure 6-B), showed considerable variation among the different regions tested. Areas with high heritability, such as those located in the southern region of Brazil, indicate higher experimental quality and the expression of genetic differences, which is crucial for the selection of superior genotypes, making these areas suitable for testing new lines. While the variation in heritability may partly reflect how the trials were conducted, rather than specific characteristics of the region, it is important to consider that lower environmental variability in these areas may also have contributed to the increased heritability. Thus, the choice of test location can influence the success of genetic selection, serving as a guide to identify environments where selection will be more efficient.

The Mega-environments (Figure 6-C) show the genetic correlation among projected values across the TPE, aiming to group pixels into regions with high genotypic correlation. However, as shown in the figure, it was not possible to define consistent macrozones for rice breeding using our dataset and methodology. The MEs were highly fragmented, and most of the territory lacked sufficient genotypic correlation to be grouped, leading to classification into a non-grouping ME zone. This fragmentation suggests that, although some areas within the same ME may show similar phenotypic responses, the lack of genetic correlation across much of Brazil limits the formation of broad and continuous mega-environments. This contrasts with the findings of Callister et al. (2024) for *Eucalyptus globulus* in Australia. Our results emphasize the need to consider not only geographic proximity but also genetic and environmental similarity when defining mega-environments to allow for more effective genotype selection—even between geographically distant locations. Similar challenges were reported by Krause et al. (2022), who observed that soybean breeding regions in North America could be grouped into mega-environments, but genotypic variation remained strongly influenced by location-specific factors.

The interpolated PA of the models (Figure 6-D) indicates the model’s accuracy in predicting phenotypic performance for the region of interest. Regions colored closer to violet tend to show real values that are closer to those predicted by the model, indicating a higher level of initial reliability. In these cases, resources can be optimized by reducing the need for real experiments in those areas, as enviromics can accurately predict crop performance. On the other hand, in regions where PA is low, it becomes more necessary to conduct real experiments to obtain more accurate data. Studies such as Jarquin et al. (2018) show that the predictive ability of models can be substantially improved by considering appropriate interactions between variables, reinforcing the importance of using efficient models to more accurately predict phenotypic performance in untested environments, thus better directing resource use within a breeding program.

The prediction accuracy of the model used in this study was dependent both on the type of variable used and on the number of covariates employed in the model. The validation process, separating training and validation groups in an 80:20 ratio, returned some interesting results. When 81 environmental covariates were reached, a peak followed by a plateau around 53% accuracy was observed. It is worth mentioning that although the peak is reached with 81 environmental covariates, it does not make sense to use only this number of covariates for our models, as we simply cannot infer with certainty which 81 of the 383 covariates will be selected to compose the model and what the optimal arrangement between them would be to form the training and validation groups. Therefore, it is valid to use the complete set of covariates, given that collecting this data is virtually cost-free, and the expected range of variation in predictive ability will be smaller. This approach aligns well with the results of Rogers and Holland (2022), who emphasize that increasing the number of predictor variables can lead to a significant improvement in model accuracy, maximizing the efficiency of phenotypic outcome prediction.

As for alternative validation methods, L1YO presented the closest result between the traditional model and the enviromic model (Table 1). This validation included the twelve genotypes with continuous data between 1995 and 2022, with data from 1995-2021 used to predict and validate those of 2022. This similarity between results is due to the trend in average productivity, represented by the average BLUP values of the genotypes, being similar over time, especially when using sequential data. When comparing the basic G×E interaction model with the model using raw environmental covariates in the other two validation processes (between quadrants and LOO), the superiority of using enviromic models is evident for our dataset. Predictive ability improved from a maximum of 8.2% in the traditional model to a minimum of 51.0% when using the model with raw environmental covariates, demonstrating the effectiveness of enviromic models in capturing environmental complexity and improving genotypic performance prediction.

## 5. Final Considerations

In this study, we demonstrated the effectiveness of using integrated environmental covariates to build prediction models to assess rice grain yield in different environments across Brazil. The analysis revealed that utilizing a comprehensive set of environmental covariates is essential for increasing prediction accuracy and better understanding genotype-environment interactions. The enviromic approach applied in this work, considering 383 environmental covariates, successfully predicted grain yield with high accuracy in certain regions using environmental data. In regions with high PA, it is possible to optimize resources by partially replacing field trials with virtual trials, allowing for cost savings that can be redirected toward conducting field trials in regions with low PA, where real experimentation is more necessary. This strategic use of virtual trials can increase the efficiency of breeding programs without compromising result accuracy.

Our results also highlighted that environmental dissimilarities are more relevant than geographical distance in explaining phenotypic response, suggesting that models using environmental covariates can provide more accurate recommendations for genotype utilization in new agricultural areas. The results showed that the attempt to define mega-environments for upland rice was not effective due to the large environmental and genotypic variation observed, and probably because the number of trials was insufficient to cover the vast Brazilian territory. This difficulty in establishing mega-environments indicates that a regionalized, locally adjusted approach, rather than broad zoning, may be more appropriate for optimizing the selection of suitable genotypes.

## Statements & Declarations

## Author Contributions

**M.A.M.B.:** Conceptualization; Methodology; Formal analysis; Data Curation; Writing - Original Draft; Visualization. **G.E.M.:** Formal analysis; Investigation; Data Curation. **F.B.:** Resources; Writing - Review & Editing. **K.O.G.D.:** Writing - Review & Editing. **P.G.S.M.:** Writing - Review & Editing. **Y.X.:** Writing - Review & Editing. **R.T.R.:** Conceptualization; Methodology; Formal analysis; Writing - Original Draft; Visualization; Supervision; Project administration. All authors reviewed the manuscript and validated the results.

## Consent to participate

Informed consent was obtained from all individual participants included in the study.

## Acknowledgments

The Graduate Program in Genetics and Plant Breeding at the Federal University of Goiás (PPGGMP/UFG), Brazilian Agricultural Research Corporation (EMBRAPA) for providing the upland rice data. Thanks to the Goiás Research Support Foundation (FAPEG) and National Council for Scientific and Technological Development (CNPQ) for providing research scholarships, and to Precision Breeding Laboratory (LAMP) for the physical infrastructure. Special thanks to student Cesar Augusto Galvão from University of Brasília (UnB) for compiling the covariates dictionary, to the Dr. Raimundo Nonato Carvalho da Rocha from Embrapa for provide the UAV image used on Figure 2, and to Professor Alexandre Coelho from Federal University of Goiás (UFG) for suggestions on statistical analyses.

