## Supplementary Material for "Predictive Ability of Enviromic Modeling in G×E Interactions for Upland Rice Site Recommendations"

### == SUPLEMENTARY MATERIALS ==

**Table S1**. List and description of environmental (or envirotypic) covariates used.

| Data Base | Cod | Acronym | Description |
|---|---|---|---|
| WorldClim[1] | X001 | bio_1 | Annual Mean Temperature |
| WorldClim | X002 | bio_2 | Mean Diurnal Range (Mean of monthly (temp max - temp min)) |
| WorldClim | X003 | bio_3 | Isothermality (bio_2/bio_7) (×100) |
| WorldClim | X004 | bio_4 | Temperature Seasonality (standard deviation * 100) |
| WorldClim | X005 | bio_5 | Max Temperature of Warmest Month |
| WorldClim | X006 | bio_6 | Min Temperature of Coldest Month |
| WorldClim | X007 | bio_7 | Temperature Annual Range (bio_5 - bio_6) |
| WorldClim | X008 | bio_8 | Mean Temperature of Wettest Quarter |
| WorldClim | X009 | bio_9 | Mean Temperature of Driest Quarter |
| WorldClim | X010 | bio_10 | Mean Temperature of Warmest Quarter |
| WorldClim | X011 | bio_11 | Mean Temperature of Coldest Quarter |
| WorldClim | X012 | bio_12 | Annual Precipitation |
| WorldClim | X013 | bio_13 | Precipitation of Wettest Month |
| WorldClim | X014 | bio_14 | Precipitation of Driest Month |
| WorldClim | X015 | bio_15 | Precipitation Seasonality (Coefficient of Variation) |
| WorldClim | X016 | bio_16 | Precipitation of Wettest Quarter |
| WorldClim | X017 | bio_17 | Precipitation of Driest Quarter |
| WorldClim | X018 | bio_18 | Precipitation of Warmest Quarter |
| WorldClim | X019 | bio_19 | Precipitation of Coldest Quarter |
| SoilGrids[2] | X020 | bdod_0-5cm_mean | Bulk Density of Fine Earth Fraction in $cg/cm^3$ for 0 to 5 cm - Mean |
| SoilGrids | X021 | bdod_0-5cm_Q0p05 | Bulk Density of Fine Earth Fraction in $cg/cm^3$ for 0 to 5 cm - Prediction for 0.05 Quantile |
| SoilGrids | X022 | bdod_0-5cm_Q0p5 | Bulk Density of Fine Earth Fraction in $cg/cm^3$ for 0 to 5 cm - Prediction for 0.5 Quantile |
| SoilGrids | X023 | bdod_0-5cm_Q0p95 | Bulk Density of Fine Earth Fraction in $cg/cm^3$ for 0 to 5 cm - Prediction for 0.95 Quantile |
| SoilGrids | X024 | bdod_5-15cm_mean | Bulk Density of Fine Earth Fraction in $cg/cm^3$ for 5 to 15 cm - Mean |
| SoilGrids | X025 | bdod_5-15cm_Q0p05 | Bulk Density of Fine Earth Fraction in $cg/cm^3$ for 5 to 15 cm - Prediction for 0.05 Quantile |
| SoilGrids | X026 | bdod_5-15cm_Q0p5 | Bulk Density of Fine Earth Fraction in $cg/cm^3$ for 5 to 15 cm - Prediction for 0.5 Quantile |
| SoilGrids | X027 | bdod_5-15cm_Q0p95 | Bulk Density of Fine Earth Fraction in $cg/cm^3$ for 5 to 15 cm - Prediction for 0.95 Quantile |
| SoilGrids | X028 | bdod_15-30cm_mean | Bulk Density of Fine Earth Fraction in $cg/cm^3$ for 15 to 30 cm - Mean |
| SoilGrids | X029 | bdod_15-30cm_Q0p05 | Bulk Density of Fine Earth Fraction in $cg/cm^3$ for 15 to 30 cm - Prediction for 0.05 Quantile |

| | | | |
|---|---|---|---|
| SoilGrids | X030 | bdod_15-30cm_Q0p5 | Bulk Density of Fine Earth Fraction in cg/cm³ for 15 to 30 cm - Prediction for 0.5 Quantile |
| SoilGrids | X031 | bdod_15-30cm_Q0p95 | Bulk Density of Fine Earth Fraction in cg/cm³ for 15 to 30 cm - Prediction for 0.95 Quantile |
| SoilGrids | X032 | bdod_30-60cm_mean | Bulk Density of Fine Earth Fraction in cg/cm³ for 30 to 60 cm - Mean |
| SoilGrids | X033 | bdod_30-60cm_Q0p05 | Bulk Density of Fine Earth Fraction in cg/cm³ for 30 to 60 cm - Prediction for 0.05 Quantile |
| SoilGrids | X034 | bdod_30-60cm_Q0p5 | Bulk Density of Fine Earth Fraction in cg/cm³ for 30 to 60 cm - Prediction for 0.5 Quantile |
| SoilGrids | X035 | bdod_30-60cm_Q0p95 | Bulk Density of Fine Earth Fraction in cg/cm³ for 30 to 60 cm - Prediction for 0.95 Quantile |
| SoilGrids | X036 | bdod_60-100cm_mean | Bulk Density of Fine Earth Fraction in cg/cm³ for 60 to 100 cm - Mean |
| SoilGrids | X037 | bdod_60-100cm_Q0p05 | Bulk Density of Fine Earth Fraction in cg/cm³ for 60 to 100 cm - Prediction for 0.05 Quantile |
| SoilGrids | X038 | bdod_60-100cm_Q0p5 | Bulk Density of Fine Earth Fraction in cg/cm³ for 60 to 100 cm - Prediction for 0.5 Quantile |
| SoilGrids | X039 | bdod_60-100cm_Q0p95 | Bulk Density of Fine Earth Fraction in cg/cm³ for 60 to 100 cm - Prediction for 0.95 Quantile |
| SoilGrids | X040 | bdod_100-200cm_mean | Bulk Density of Fine Earth Fraction in cg/cm³ for 100 to 200 cm - Mean |
| SoilGrids | X041 | bdod_100-200cm_Q0p05 | Bulk Density of Fine Earth Fraction in cg/cm³ for 100 to 200 cm - Prediction for 0.05 Quantile |
| SoilGrids | X042 | bdod_100-200cm_Q0p5 | Bulk Density of Fine Earth Fraction in cg/cm³ for 100 to 200 cm - Prediction for 0.5 Quantile |
| SoilGrids | X043 | bdod_100-200cm_Q0p95 | Bulk Density of Fine Earth Fraction in cg/cm³ for 100 to 200 cm - Prediction for 0.95 Quantile |
| SoilGrids | X044 | cec_0-5cm_mean | Cation Exchange Capacity (CEC) in mmol(c)/kg for 0 to 5 cm - Mean |
| SoilGrids | X045 | cec_0-5cm_Q0p05 | Cation Exchange Capacity (CEC) in mmol(c)/kg for 0 to 5 cm - Prediction for 0.05 Quantile |
| SoilGrids | X046 | cec_0-5cm_Q0p5 | Cation Exchange Capacity (CEC) in mmol(c)/kg for 0 to 5 cm - Prediction for 0.5 Quantile |
| SoilGrids | X047 | cec_0-5cm_Q0p95 | Cation Exchange Capacity (CEC) in mmol(c)/kg for 0 to 5 cm - Prediction for 0.95 Quantile |
| SoilGrids | X048 | cec_5-15cm_mean | Cation Exchange Capacity (CEC) in mmol(c)/kg for 5 to 15 cm - Mean |
| SoilGrids | X049 | cec_5-15cm_Q0p05 | Cation Exchange Capacity (CEC) in mmol(c)/kg for 5 to 15 cm - Prediction for 0.05 Quantile |
| SoilGrids | X050 | cec_5-15cm_Q0p5 | Cation Exchange Capacity (CEC) in mmol(c)/kg for 5 to 15 cm - Prediction for 0.5 Quantile |
| SoilGrids | X051 | cec_5-15cm_Q0p95 | Cation Exchange Capacity (CEC) in mmol(c)/kg for 5 to 15 cm - Prediction for 0.95 Quantile |
| SoilGrids | X052 | cec_15-30cm_mean | Cation Exchange Capacity (CEC) in mmol(c)/kg for 15 to 30 cm - Mean |
| SoilGrids | X053 | cec_15-30cm_Q0p05 | Cation Exchange Capacity (CEC) in mmol(c)/kg for 15 to 30 cm - Prediction for 0.05 Quantile |
| SoilGrids | X054 | cec_15-30cm_Q0p5 | Cation Exchange Capacity (CEC) in mmol(c)/kg for 15 to 30 cm - Prediction for 0.5 Quantile |

| | | | |
|---|---|---|---|
| SoilGrids | X055 | cec_15-30cm_Q0p95 | Cation Exchange Capacity (CEC) in mmol(c)/kg for 15 to 30 cm - Prediction for 0.95 Quantile |
| SoilGrids | X056 | cec_30-60cm_mean | Cation Exchange Capacity (CEC) in mmol(c)/kg for 30 to 60 cm - Mean |
| SoilGrids | X057 | cec_30-60cm_Q0p05 | Cation Exchange Capacity (CEC) in mmol(c)/kg for 30 to 60 cm - Prediction for 0.05 Quantile |
| SoilGrids | X058 | cec_30-60cm_Q0p5 | Cation Exchange Capacity (CEC) in mmol(c)/kg for 30 to 60 cm - Prediction for 0.5 Quantile |
| SoilGrids | X059 | cec_30-60cm_Q0p95 | Cation Exchange Capacity (CEC) in mmol(c)/kg for 30 to 60 cm - Prediction for 0.95 Quantile |
| SoilGrids | X060 | cec_60-100cm_mean | Cation Exchange Capacity (CEC) in mmol(c)/kg for 60 to 100 cm - Mean |
| SoilGrids | X061 | cec_60-100cm_Q0p05 | Cation Exchange Capacity (CEC) in mmol(c)/kg for 60 to 100 cm - Prediction for 0.05 Quantile |
| SoilGrids | X062 | cec_60-100cm_Q0p5 | Cation Exchange Capacity (CEC) in mmol(c)/kg for 60 to 100 cm - Prediction for 0.5 Quantile |
| SoilGrids | X063 | cec_60-100cm_Q0p95 | Cation Exchange Capacity (CEC) in mmol(c)/kg for 60 to 100 cm - Prediction for 0.95 Quantile |
| SoilGrids | X064 | cec_100-200cm_mean | Cation Exchange Capacity (CEC) in mmol(c)/kg for 100 to 200 cm - Mean |
| SoilGrids | X065 | cec_100-200cm_Q0p05 | Cation Exchange Capacity (CEC) in mmol(c)/kg for 100 to 200 cm - Prediction for 0.05 Quantile |
| SoilGrids | X066 | cec_100-200cm_Q0p5 | Cation Exchange Capacity (CEC) in mmol(c)/kg for 100 to 200 cm - Prediction for 0.5 Quantile |
| SoilGrids | X067 | cec_100-200cm_Q0p95 | Cation Exchange Capacity (CEC) in mmol(c)/kg for 100 to 200 cm - Prediction for 0.95 Quantile |
| SoilGrids | X068 | cfvo_0-5cm_mean | Volumetric fraction of coarse fragments (>2 mm) in cm3/dm3 (vol‰) for 0 to 5 cm of the surface - mean |
| SoilGrids | X069 | cfvo_0-5cm_Q0p05 | Volumetric fraction of coarse fragments (>2 mm) in cm3/dm3 (vol‰) for 0 to 5 cm of the surface – prediction for quantile 0.05 |
| SoilGrids | X070 | cfvo_0-5cm_Q0p5 | Volumetric fraction of coarse fragments (>2 mm) in cm3/dm3 (vol‰) for 0 to 5 cm of the surface – prediction for quantile 0.5 |
| SoilGrids | X071 | cfvo_0-5cm_Q0p95 | Volumetric fraction of coarse fragments (>2 mm) in cm3/dm3 (vol‰) for 0 to 5 cm of the surface – prediction for quantile 0.95 |
| SoilGrids | X072 | cfvo_5-15cm_mean | Volumetric fraction of coarse fragments (>2 mm) in cm3/dm3 (vol‰) for 5 to 15 cm of the surface - mean |
| SoilGrids | X073 | cfvo_5-15cm_Q0p05 | Volumetric fraction of coarse fragments (>2 mm) in cm3/dm3 (vol‰) for 5 to 15 cm of the surface – prediction for quantile 0.05 |
| SoilGrids | X074 | cfvo_5-15cm_Q0p5 | Volumetric fraction of coarse fragments (>2 mm) in cm3/dm3 (vol‰) for 5 to 15 cm of the surface – prediction for quantile 0.5 |
| SoilGrids | X075 | cfvo_5-15cm_Q0p95 | Volumetric fraction of coarse fragments (>2 mm) in cm3/dm3 (vol‰) for 5 to 15 cm of the surface – prediction for quantile 0.95 |
| SoilGrids | X076 | cfvo_15-30cm_mean | Volumetric fraction of coarse fragments (>2 mm) in cm3/dm3 (vol‰) for 15 to 30 cm of the surface - mean |

| | | | |
|---|---|---|---|
| SoilGrids | X077 | cfvo_15-30cm_Q0p05 | Volumetric fraction of coarse fragments (>2 mm) in cm3/dm3 (vol‰) for 15 to 30 cm of the surface – prediction for quantile 0.05 |
| SoilGrids | X078 | cfvo_15-30cm_Q0p5 | Volumetric fraction of coarse fragments (>2 mm) in cm3/dm3 (vol‰) for 15 to 30 cm of the surface – prediction for quantile 0.5 |
| SoilGrids | X079 | cfvo_15-30cm_Q0p95 | Volumetric fraction of coarse fragments (>2 mm) in cm3/dm3 (vol‰) for 15 to 30 cm of the surface – prediction for quantile 0.95 |
| SoilGrids | X080 | cfvo_30-60cm_mean | Volumetric fraction of coarse fragments (>2 mm) in cm3/dm3 (vol‰) for 30 to 60 cm of the surface - mean |
| SoilGrids | X081 | cfvo_30-60cm_Q0p05 | Volumetric fraction of coarse fragments (>2 mm) in cm3/dm3 (vol‰) for 30 to 60 cm of the surface – prediction for quantile 0.05 |
| SoilGrids | X082 | cfvo_30-60cm_Q0p5 | Volumetric fraction of coarse fragments (>2 mm) in cm3/dm3 (vol‰) for 30 to 60 cm of the surface – prediction for quantile 0.5 |
| SoilGrids | X083 | cfvo_30-60cm_Q0p95 | Volumetric fraction of coarse fragments (>2 mm) in cm3/dm3 (vol‰) for 30 to 60 cm of the surface – prediction for quantile 0.95 |
| SoilGrids | X084 | cfvo_60-100cm_mean | Volumetric fraction of coarse fragments (>2 mm) in cm3/dm3 (vol‰) for 60 to 100 cm of the surface - mean |
| SoilGrids | X085 | cfvo_60-100cm_Q0p05 | Volumetric fraction of coarse fragments (>2 mm) in cm3/dm3 (vol‰) for 60 to 100 cm of the surface – prediction for quantile 0.05 |
| SoilGrids | X086 | cfvo_60-100cm_Q0p5 | Volumetric fraction of coarse fragments (>2 mm) in cm3/dm3 (vol‰) for 60 to 100 cm of the surface – prediction for quantile 0.5 |
| SoilGrids | X087 | cfvo_60-100cm_Q0p95 | Volumetric fraction of coarse fragments (>2 mm) in cm3/dm3 (vol‰) for 60 to 100 cm of the surface – prediction for quantile 0.95 |
| SoilGrids | X088 | cfvo_100-200cm_mean | Volumetric fraction of coarse fragments (>2 mm) in cm3/dm3 (vol‰) for 100 to 200 cm of the surface - mean |
| SoilGrids | X089 | cfvo_100-200cm_Q0p05 | Volumetric fraction of coarse fragments (>2 mm) in cm3/dm3 (vol‰) for 100 to 200 cm of the surface – prediction for quantile 0.05 |
| SoilGrids | X090 | cfvo_100-200cm_Q0p5 | Volumetric fraction of coarse fragments (>2 mm) in cm3/dm3 (vol‰) for 100 to 200 cm of the surface – prediction for quantile 0.5 |
| SoilGrids | X091 | cfvo_100-200cm_Q0p95 | Volumetric fraction of coarse fragments (>2 mm) in cm3/dm3 (vol‰) for 100 to 200 cm of the surface – prediction for quantile 0.95 |
| SoilGrids | X092 | clay_0-5cm_mean | Proportion of clay particles (<0.002 mm) in fine soil fraction in g/kg for 0 to 5 cm of the surface - mean |
| SoilGrids | X093 | clay_0-5cm_Q0p05 | Proportion of clay particles (<0.002 mm) in fine soil fraction in g/kg for 0 to 5 cm of the surface – prediction for quantile 0.05 |
| SoilGrids | X094 | clay_0-5cm_Q0p5 | Proportion of clay particles (<0.002 mm) in fine soil fraction in g/kg for 0 to 5 cm of the surface – prediction for quantile 0.5 |

| | | | |
|---|---|---|---|
| SoilGrids | X095 | clay_0-5cm_Q0p95 | Proportion of clay particles (<0.002 mm) in fine soil fraction in g/kg for 0 to 5 cm of the surface – prediction for quantile 0.95 |
| SoilGrids | X096 | clay_5-15cm_mean | Proportion of clay particles (<0.002 mm) in fine soil fraction in g/kg for 5 to 15 cm of the surface - mean |
| SoilGrids | X097 | clay_5-15cm_Q0p05 | Proportion of clay particles (<0.002 mm) in fine soil fraction in g/kg for 5 to 15 cm of the surface – prediction for quantile 0.05 |
| SoilGrids | X098 | clay_5-15cm_Q0p5 | Proportion of clay particles (<0.002 mm) in fine soil fraction in g/kg for 5 to 15 cm of the surface – prediction for quantile 0.5 |
| SoilGrids | X099 | clay_5-15cm_Q0p95 | Proportion of clay particles (<0.002 mm) in fine soil fraction in g/kg for 5 to 15 cm of the surface – prediction for quantile 0.95 |
| SoilGrids | X100 | clay_15-30cm_mean | Proportion of clay particles (<0.002 mm) in fine soil fraction in g/kg for 15 to 30 cm of the surface - mean |
| SoilGrids | X101 | clay_15-30cm_Q0p05 | Proportion of clay particles (<0.002 mm) in fine soil fraction in g/kg for 15 to 30 cm of the surface – prediction for quantile 0.05 |
| SoilGrids | X102 | clay_15-30cm_Q0p5 | Proportion of clay particles (<0.002 mm) in fine soil fraction in g/kg for 15 to 30 cm of the surface – prediction for quantile 0.5 |
| SoilGrids | X103 | clay_15-30cm_Q0p95 | Proportion of clay particles (<0.002 mm) in fine soil fraction in g/kg for 15 to 30 cm of the surface – prediction for quantile 0.95 |
| SoilGrids | X104 | clay_30-60cm_mean | Proportion of clay particles (<0.002 mm) in fine soil fraction in g/kg for 30 to 60 cm of the surface - mean |
| SoilGrids | X105 | clay_30-60cm_Q0p05 | Proportion of clay particles (<0.002 mm) in fine soil fraction in g/kg for 30 to 60 cm of the surface – prediction for quantile 0.05 |
| SoilGrids | X106 | clay_30-60cm_Q0p5 | Proportion of clay particles (<0.002 mm) in fine soil fraction in g/kg for 30 to 60 cm of the surface – prediction for quantile 0.5 |
| SoilGrids | X107 | clay_30-60cm_Q0p95 | Proportion of clay particles (<0.002 mm) in fine soil fraction in g/kg for 30 to 60 cm of the surface – prediction for quantile 0.95 |
| SoilGrids | X108 | clay_60-100cm_mean | Proportion of clay particles (<0.002 mm) in fine soil fraction in g/kg for 60 to 100 cm of the surface - mean |
| SoilGrids | X109 | clay_60-100cm_Q0p05 | Proportion of clay particles (<0.002 mm) in fine soil fraction in g/kg for 60 to 100 cm of the surface – prediction for quantile 0.05 |
| SoilGrids | X110 | clay_60-100cm_Q0p5 | Proportion of clay particles (<0.002 mm) in fine soil fraction in g/kg for 60 to 100 cm of the surface – prediction for quantile 0.5 |
| SoilGrids | X111 | clay_60-100cm_Q0p95 | Proportion of clay particles (<0.002 mm) in fine soil fraction in g/kg for 60 to 100 cm of the surface – prediction for quantile 0.95 |
| SoilGrids | X112 | clay_100-200cm_mean | Proportion of clay particles (<0.002 mm) in fine soil fraction in g/kg for 100 to 200 cm of the surface - mean |

| | | | |
|---|---|---|---|
| SoilGrids | X113 | clay_100-200cm_Q0p05 | Proportion of clay particles (<0.002 mm) in fine soil fraction in g/kg for 100 to 200 cm of the surface – prediction for quantile 0.05 |
| SoilGrids | X114 | clay_100-200cm_Q0p5 | Proportion of clay particles (<0.002 mm) in fine soil fraction in g/kg for 100 to 200 cm of the surface – prediction for quantile 0.5 |
| SoilGrids | X115 | clay_100-200cm_Q0p95 | Proportion of clay particles (<0.002 mm) in fine soil fraction in g/kg for 100 to 200 cm of the surface – prediction for quantile 0.95 |
| SoilGrids | X116 | nitrogen_0- 5cm_mean | Total Nitrogen (N) cg/kg for 0 to 5 cm of the surface - mean |
| SoilGrids | X117 | nitrogen_0- 5cm_Q0p05 | Total Nitrogen (N) cg/kg for 0 to 5 cm of the surface – prediction for quantile 0.05 |
| SoilGrids | X118 | nitrogen_0- 5cm_Q0p5 | Total Nitrogen (N) cg/kg for 0 to 5 cm of the surface – prediction for quantile 0.5 |
| SoilGrids | X119 | nitrogen_0- 5cm_Q0p95 | Total Nitrogen (N) cg/kg for 0 to 5 cm of the surface – prediction for quantile 0.95 |
| SoilGrids | X120 | nitrogen_5-15cm_mean | Total Nitrogen (N) cg/kg for 5 to 15 cm of the surface - mean |
| SoilGrids | X121 | nitrogen_5-15cm_Q0p05 | Total Nitrogen (N) cg/kg for 5 to 15 cm of the surface – prediction for quantile 0.05 |
| SoilGrids | X122 | nitrogen_5-15cm_Q0p5 | Total Nitrogen (N) cg/kg for 5 to 15 cm of the surface – prediction for quantile 0.5 |
| SoilGrids | X123 | nitrogen_5-15cm_Q0p95 | Total Nitrogen (N) cg/kg for 5 to 15 cm of the surface – prediction for quantile 0.95 |
| SoilGrids | X124 | nitrogen_15-30cm_mean | Total Nitrogen (N) cg/kg for 15 to 30 cm of the surface - mean |
| SoilGrids | X125 | nitrogen_15-30cm_Q0p05 | Total Nitrogen (N) cg/kg for 15 to 30 cm of the surface – prediction for quantile 0.05 |
| SoilGrids | X126 | nitrogen_15-30cm_Q0p5 | Total Nitrogen (N) cg/kg for 15 to 30 cm of the surface – prediction for quantile 0.5 |
| SoilGrids | X127 | nitrogen_15-30cm_Q0p95 | Total Nitrogen (N) cg/kg for 15 to 30 cm of the surface – prediction for quantile 0.95 |
| SoilGrids | X128 | nitrogen_30-60cm_mean | Total Nitrogen (N) cg/kg for 30 to 60 cm of the surface - mean |
| SoilGrids | X129 | nitrogen_30-60cm_Q0p05 | Total Nitrogen (N) cg/kg for 30 to 60 cm of the surface – prediction for quantile 0.05 |
| SoilGrids | X130 | nitrogen_30-60cm_Q0p5 | Total Nitrogen (N) cg/kg for 30 to 60 cm of the surface – prediction for quantile 0.5 |
| SoilGrids | X131 | nitrogen_30-60cm_Q0p95 | Total Nitrogen (N) cg/kg for 30 to 60 cm of the surface – prediction for quantile 0.95 |
| SoilGrids | X132 | nitrogen_60-100cm_mean | Total Nitrogen (N) cg/kg for 60 to 100 cm of the surface - mean |
| SoilGrids | X133 | nitrogen_60-100cm_Q0p05 | Total Nitrogen (N) cg/kg for 60 to 100 cm of the surface – prediction for quantile 0.05 |
| SoilGrids | X134 | nitrogen_60-100cm_Q0p5 | Total Nitrogen (N) cg/kg for 60 to 100 cm of the surface – prediction for quantile 0.5 |
| SoilGrids | X135 | nitrogen_60-1005cm_Q0p95 | Total Nitrogen (N) cg/kg for 60 to 100 cm of the surface – prediction for quantile 0.95 |
| SoilGrids | X136 | nitrogen_100-200cm_mean | Total Nitrogen (N) cg/kg for 100 to 200 cm of the surface - mean |

| | | | |
|---|---|---|---|
| SoilGrids | X137 | nitrogen_100-200cm_Q0p05 | Total Nitrogen (N) cg/kg for 100 to 200 cm of the surface – prediction for quantile 0.05 |
| SoilGrids | X138 | nitrogen_100-200cm_Q0p5 | Total Nitrogen (N) cg/kg for 100 to 200 cm of the surface – prediction for quantile 0.5 |
| SoilGrids | X139 | nitrogen_100-200cm_Q0p95 | Total Nitrogen (N) cg/kg for 100 to 200 cm of the surface – prediction for quantile 0.95 |
| SoilGrids | X140 | ocd_0-5cm_mean | Organic Carbon Density in hg/m³ for 0 to 5 cm of the surface - mean |
| SoilGrids | X141 | ocd_0-5cm_Q0p05 | Organic Carbon Density in hg/m³ for 0 to 5 cm of the surface – prediction for quantile 0.05 |
| SoilGrids | X142 | ocd_0-5cm_Q0p5 | Organic Carbon Density in hg/m³ for 0 to 5 cm of the surface – prediction for quantile 0.5 |
| SoilGrids | X143 | ocd_0-5cm_Q0p95 | Organic Carbon Density in hg/m³ for 0 to 5 cm of the surface – prediction for quantile 0.95 |
| SoilGrids | X144 | ocd_5-15cm_mean | Organic Carbon Density in hg/m³ for 5 to 15 cm of the surface - mean |
| SoilGrids | X145 | ocd_5-15cm_Q0p05 | Organic Carbon Density in hg/m³ for 5 to 15 cm of the surface – prediction for quantile 0.05 |
| SoilGrids | X146 | ocd_5-15cm_Q0p5 | Organic Carbon Density in hg/m³ for 5 to 15 cm of the surface – prediction for quantile 0.5 |
| SoilGrids | X147 | ocd_5-15cm_Q0p95 | Organic Carbon Density in hg/m³ for 5 to 15 cm of the surface – prediction for quantile 0.95 |
| SoilGrids | X148 | ocd_15-30cm_mean | Organic Carbon Density in hg/m³ for 15 to 30 cm of the surface - mean |
| SoilGrids | X149 | ocd_15-30cm_Q0p05 | Organic Carbon Density in hg/m³ for 15 to 30 cm of the surface – prediction for quantile 0.05 |
| SoilGrids | X150 | ocd_15-30cm_Q0p5 | Organic Carbon Density in hg/m³ for 15 to 30 cm of the surface – prediction for quantile 0.5 |
| SoilGrids | X151 | ocd_15-30cm_Q0p95 | Organic Carbon Density in hg/m³ for 15 to 30 cm of the surface – prediction for quantile 0.95 |
| SoilGrids | X152 | ocd_30-60cm_mean | Organic Carbon Density in hg/m³ for 30 to 60 cm of the surface - mean |
| SoilGrids | X153 | ocd_30-60cm_Q0p05 | Organic Carbon Density in hg/m³ for 30 to 60 cm of the surface – prediction for quantile 0.05 |
| SoilGrids | X154 | ocd_30-60cm_Q0p5 | Organic Carbon Density in hg/m³ for 30 to 60 cm of the surface – prediction for quantile 0.5 |
| SoilGrids | X155 | ocd_30-60cm_Q0p95 | Organic Carbon Density in hg/m³ for 30 to 60 cm of the surface – prediction for quantile 0.95 |
| SoilGrids | X156 | ocd_60-100cm_mean | Organic Carbon Density in hg/m³ for 60 to 100 cm of the surface - mean |
| SoilGrids | X157 | ocd_60-100cm_Q0p05 | Organic Carbon Density in hg/m³ for 60 to 100 cm of the surface – prediction for quantile 0.05 |
| SoilGrids | X158 | ocd_60-100cm_Q0p5 | Organic Carbon Density in hg/m³ for 60 to 100 cm of the surface – prediction for quantile 0.5 |
| SoilGrids | X159 | ocd_60-100cm_Q0p95 | Organic Carbon Density in hg/m³ for 60 to 100 cm of the surface – prediction for quantile 0.95 |
| SoilGrids | X160 | ocd_100-200cm_mean | Organic Carbon Density in hg/m³ for 100 to 200 cm of the surface - mean |
| SoilGrids | X161 | ocd_100-200cm_Q0p05 | Organic Carbon Density in hg/m³ for 100 to 200 cm of the surface – prediction for quantile 0.05 |

| | | | |
|---|---|---|---|
| SoilGrids | X162 | ocd_100-200cm_Q0p5 | Organic Carbon Density in hg/m³ for 100 to 200 cm of the surface – prediction for quantile 0.5 |
| SoilGrids | X163 | ocd_100-200cm_Q0p95 | Organic Carbon Density in hg/m³ for 100 to 200 cm of the surface – prediction for quantile 0.95 |
| SoilGrids | X164 | ocs_0-30cm_mean | Organic carbon stocks in t/ha for 0 to 30 cm of the surface - mean |
| SoilGrids | X165 | ocs_0-30cm_Q0p05 | Organic carbon stocks in t/ha for 0 to 30 cm of the surface – prediction for quantile 0.05 |
| SoilGrids | X166 | ocs_0-30cm_Q0p5 | Organic carbon stocks in t/ha for 0 to 30 cm of the surface – prediction for quantile 0.5 |
| SoilGrids | X167 | ocs_0-30cm_Q0p95 | Organic carbon stocks in t/ha for 0 to 30 cm of the surface – prediction for quantile 0.95 |
| SoilGrids | X168 | phh2o_0-5cm_mean | Soil pH in pHx10 for 0 to 5 cm of the surface - mean |
| SoilGrids | X169 | phh2o_0-5cm_Q0p05 | Soil pH in pHx10 for 0 to 5 cm of the surface – prediction for quantile 0.05 |
| SoilGrids | X170 | phh2o_0-5cm_Q0p5 | Soil pH in pHx10 for 0 to 5 cm of the surface – prediction for quantile 0.5 |
| SoilGrids | X171 | phh2o_0-5cm_Q0p95 | Soil pH in pHx10 for 0 to 5 cm of the surface – prediction for quantile 0.95 |
| SoilGrids | X172 | phh2o_5-15cm_mean | Soil pH in pHx10 for 5 to 15 cm of the surface - mean |
| SoilGrids | X173 | phh2o_5-15cm_Q0p05 | Soil pH in pHx10 for 5 to 15 cm of the surface – prediction for quantile 0.05 |
| SoilGrids | X174 | phh2o_5-15cm_Q0p5 | Soil pH in pHx10 for 5 to 15 cm of the surface – prediction for quantile 0.5 |
| SoilGrids | X175 | phh2o_5-15cm_Q0p95 | Soil pH in pHx10 for 5 to 15 cm of the surface – prediction for quantile 0.95 |
| SoilGrids | X176 | phh2o_15-30cm_mean | Soil pH in pHx10 for 15 to 30 cm of the surface - mean |
| SoilGrids | X177 | phh2o_15-30cm_Q0p05 | Soil pH in pHx10 for 15 to 30 cm of the surface – prediction for quantile 0.05 |
| SoilGrids | X178 | phh2o_15-30cm_Q0p5 | Soil pH in pHx10 for 15 to 30 cm of the surface – prediction for quantile 0.5 |
| SoilGrids | X179 | phh2o_15-30cm_Q0p95 | Soil pH in pHx10 for 15 to 30 cm of the surface – prediction for quantile 0.95 |
| SoilGrids | X180 | phh2o_30-60cm_mean | Soil pH in pHx10 for 30 to 60 cm of the surface - mean |
| SoilGrids | X181 | phh2o_30-60cm_Q0p05 | Soil pH in pHx10 for 30 to 60 cm of the surface – prediction for quantile 0.05 |
| SoilGrids | X182 | phh2o_30-60cm_Q0p5 | Soil pH in pHx10 for 30 to 60 cm of the surface – prediction for quantile 0.5 |
| SoilGrids | X183 | phh2o_30-60cm_Q0p95 | Soil pH in pHx10 for 30 to 60 cm of the surface – prediction for quantile 0.95 |
| SoilGrids | X184 | phh2o_60-100cm_mean | Soil pH in pHx10 for 60 to 100 cm of the surface - mean |
| SoilGrids | X185 | phh2o_60-100cm_Q0p05 | Soil pH in pHx10 for 60 to 100 cm of the surface – prediction for quantile 0.05 |
| SoilGrids | X186 | phh2o_60-100cm_Q0p5 | Soil pH in pHx10 for 60 to 100 cm of the surface – prediction for quantile 0.5 |
| SoilGrids | X187 | phh2o_60-100cm_Q0p95 | Soil pH in pHx10 for 60 to 100 cm of the surface – prediction for quantile 0.95 |
| SoilGrids | X188 | phh2o_100-200cm_mean | Soil pH in pHx10 for 100 to 200 cm of the surface - mean |

| | | | |
|---|---|---|---|
| SoilGrids | X189 | phh2o_100-200cm_Q0p05 | Soil pH in pHx10 for 100 to 200 cm of the surface – prediction for quantile 0.05 |
| SoilGrids | X190 | phh2o_100-200cm_Q0p5 | Soil pH in pHx10 for 100 to 200 cm of the surface – prediction for quantile 0.5 |
| SoilGrids | X191 | phh2o_100-200cm_Q0p95 | Soil pH in pHx10 for 100 to 200 cm of the surface – prediction for quantile 0.95 |
| SoilGrids | X192 | sand_0-5cm_mean | Sand particle proportion (>0.05 mm) for fine soil fraction in g/kg for 0 to 5 cm of the surface - mean |
| SoilGrids | X193 | sand_0-5cm_Q0p05 | Sand particle proportion (>0.05 mm) for fine soil fraction in g/kg for 0 to 5 cm of the surface – prediction for quantile 0.05 |
| SoilGrids | X194 | sand_0-5cm_Q0p5 | Sand particle proportion (>0.05 mm) for fine soil fraction in g/kg for 0 to 5 cm of the surface – prediction for quantile 0.5 |
| SoilGrids | X195 | sand_0-5cm_Q0p95 | Sand particle proportion (>0.05 mm) for fine soil fraction in g/kg for 0 to 5 cm of the surface – prediction for quantile 0.95 |
| SoilGrids | X196 | sand_5-15cm_mean | Sand particle proportion (>0.05 mm) for fine soil fraction in g/kg for 5 to 15 cm of the surface - mean |
| SoilGrids | X197 | sand_5-15cm_Q0p05 | Sand particle proportion (>0.05 mm) for fine soil fraction in g/kg for 5 to 15 cm of the surface – prediction for quantile 0.05 |
| SoilGrids | X198 | sand_5-15cm_Q0p5 | Sand particle proportion (>0.05 mm) for fine soil fraction in g/kg for 5 to 15 cm of the surface – prediction for quantile 0.5 |
| SoilGrids | X199 | sand_5-15cm_Q0p95 | Sand particle proportion (>0.05 mm) for fine soil fraction in g/kg for 5 to 15 cm of the surface – prediction for quantile 0.95 |
| SoilGrids | X200 | sand_15-30cm_mean | Sand particle proportion (>0.05 mm) for fine soil fraction in g/kg for 15 to 30 cm of the surface - mean |
| SoilGrids | X201 | sand_15-30cm_Q0p05 | Sand particle proportion (>0.05 mm) for fine soil fraction in g/kg for 15 to 30 cm of the surface – prediction for quantile 0.05 |
| SoilGrids | X202 | sand_15-30cm_Q0p5 | Sand particle proportion (>0.05 mm) for fine soil fraction in g/kg for 15 to 30 cm of the surface – prediction for quantile 0.5 |
| SoilGrids | X203 | sand_15-30cm_Q0p95 | Sand particle proportion (>0.05 mm) for fine soil fraction in g/kg for 15 to 30 cm of the surface – prediction for quantile 0.95 |
| SoilGrids | X204 | sand_30-60cm_mean | Sand particle proportion (>0.05 mm) for fine soil fraction in g/kg for 30 to 60 cm of the surface - mean |
| SoilGrids | X205 | sand_30-60cm_Q0p05 | Sand particle proportion (>0.05 mm) for fine soil fraction in g/kg for 30 to 60 cm of the surface – prediction for quantile 0.05 |
| SoilGrids | X206 | sand_30-60cm_Q0p5 | Sand particle proportion (>0.05 mm) for fine soil fraction in g/kg for 30 to 60 cm of the surface – prediction for quantile 0.5 |
| SoilGrids | X207 | sand_30-60cm_Q0p95 | Sand particle proportion (>0.05 mm) for fine soil fraction in g/kg for 30 to 60 cm of the surface – prediction for quantile 0.95 |

| | | | |
|---|---|---|---|
| SoilGrids | X208 | sand_60-100cm_mean | Sand particle proportion (>0.05 mm) for fine soil fraction in g/kg for 60 to 100 cm of the surface - mean |
| SoilGrids | X209 | sand_60-100cm_Q0p05 | Sand particle proportion (>0.05 mm) for fine soil fraction in g/kg for 60 to 100 cm of the surface – prediction for quantile 0.05 |
| SoilGrids | X210 | sand_60-100cm_Q0p5 | Sand particle proportion (>0.05 mm) for fine soil fraction in g/kg for 60 to 100 cm of the surface – prediction for quantile 0.5 |
| SoilGrids | X211 | sand_60-100cm_Q0p95 | Sand particle proportion (>0.05 mm) for fine soil fraction in g/kg for 60 to 100 cm of the surface – prediction for quantile 0.95 |
| SoilGrids | X212 | sand_100-200cm_mean | Sand particle proportion (>0.05 mm) for fine soil fraction in g/kg for 100 to 200 cm of the surface - mean |
| SoilGrids | X213 | sand_100-200cm_Q0p05 | Sand particle proportion (>0.05 mm) for fine soil fraction in g/kg for 100 to 200 cm of the surface – prediction for quantile 0.05 |
| SoilGrids | X214 | sand_100-200cm_Q0p5 | Sand particle proportion (>0.05 mm) for fine soil fraction in g/kg for 100 to 200 cm of the surface – prediction for quantile 0.5 |
| SoilGrids | X215 | sand_100-200cm_Q0p95 | Sand particle proportion (>0.05 mm) for fine soil fraction in g/kg for 100 to 200 cm of the surface – prediction for quantile 0.95 |
| SoilGrids | X216 | silt_0-5cm_mean | Silt particle proportion (>0.05 mm) in fine soil fraction in g/kg for 0 to 5 cm of the surface - mean |
| SoilGrids | X217 | silt_0-5cm_Q0p05 | Silt particle proportion (>0.05 mm) in fine soil fraction in g/kg for 0 to 5 cm of the surface – prediction for quantile 0.05 |
| SoilGrids | X218 | silt_0-5cm_Q0p5 | Silt particle proportion (>0.05 mm) in fine soil fraction in g/kg for 0 to 5 cm of the surface – prediction for quantile 0.5 |
| SoilGrids | X219 | silt_0-5cm_Q0p95 | Silt particle proportion (>0.05 mm) in fine soil fraction in g/kg for 0 to 5 cm of the surface – prediction for quantile 0.95 |
| SoilGrids | X220 | silt_5-15cm_mean | Silt particle proportion (>0.05 mm) in fine soil fraction in g/kg for 5 to 15 cm of the surface - mean |
| SoilGrids | X221 | silt_5-15cm_Q0p05 | Silt particle proportion (>0.05 mm) in fine soil fraction in g/kg for 5 to 15 cm of the surface – prediction for quantile 0.05 |
| SoilGrids | X222 | silt_5-15cm_Q0p5 | Silt particle proportion (>0.05 mm) in fine soil fraction in g/kg for 5 to 15 cm of the surface – prediction for quantile 0.5 |
| SoilGrids | X223 | silt_5-15cm_Q0p95 | Silt particle proportion (>0.05 mm) in fine soil fraction in g/kg for 5 to 15 cm of the surface – prediction for quantile 0.95 |
| SoilGrids | X224 | silt_15-30cm_mean | Silt particle proportion (>0.05 mm) in fine soil fraction in g/kg for 15 to 30 cm of the surface - mean |
| SoilGrids | X225 | silt_15-30cm_Q0p05 | Silt particle proportion (>0.05 mm) in fine soil fraction in g/kg for 15 to 30 cm of the surface – prediction for quantile 0.05 |

| | | | |
|---|---|---|---|
| SoilGrids | X226 | silt_15-30cm_Q0p5 | Silt particle proportion (>0.05 mm) in fine soil fraction in g/kg for 15 to 30 cm of the surface – prediction for quantile 0.5 |
| SoilGrids | X227 | silt_15-30cm_Q0p95 | Silt particle proportion (>0.05 mm) in fine soil fraction in g/kg for 15 to 30 cm of the surface – prediction for quantile 0.95 |
| SoilGrids | X228 | silt_30-60cm_mean | Silt particle proportion (>0.05 mm) in fine soil fraction in g/kg for 30 to 60 cm of the surface - mean |
| SoilGrids | X229 | silt_30-60cm_Q0p05 | Silt particle proportion (>0.05 mm) in fine soil fraction in g/kg for 30 to 60 cm of the surface – prediction for quantile 0.05 |
| SoilGrids | X230 | silt_30-60cm_Q0p5 | Silt particle proportion (>0.05 mm) in fine soil fraction in g/kg for 30 to 60 cm of the surface – prediction for quantile 0.5 |
| SoilGrids | X231 | silt_30-60cm_Q0p95 | Silt particle proportion (>0.05 mm) in fine soil fraction in g/kg for 30 to 60 cm of the surface – prediction for quantile 0.95 |
| SoilGrids | X232 | silt_60-100cm_mean | Silt particle proportion (>0.05 mm) in fine soil fraction in g/kg for 60 to 100 cm of the surface - mean |
| SoilGrids | X233 | silt_60-100cm_Q0p05 | Silt particle proportion (>0.05 mm) in fine soil fraction in g/kg for 60 to 100 cm of the surface – prediction for quantile 0.05 |
| SoilGrids | X234 | silt_60-100cm_Q0p5 | Silt particle proportion (>0.05 mm) in fine soil fraction in g/kg for 60 to 100 cm of the surface – prediction for quantile 0.5 |
| SoilGrids | X235 | silt_60-100cm_Q0p95 | Silt particle proportion (>0.05 mm) in fine soil fraction in g/kg for 60 to 100 cm of the surface – prediction for quantile 0.95 |
| SoilGrids | X236 | silt_100-200cm_mean | Silt particle proportion (>0.05 mm) in fine soil fraction in g/kg for 100 to 200 cm of the surface - mean |
| SoilGrids | X237 | silt_100-200cm_Q0p05 | Silt particle proportion (>0.05 mm) in fine soil fraction in g/kg for 100 to 200 cm of the surface – prediction for quantile 0.05 |
| SoilGrids | X238 | silt_100-200cm_Q0p5 | Silt particle proportion (>0.05 mm) in fine soil fraction in g/kg for 100 to 200 cm of the surface – prediction for quantile 0.5 |
| SoilGrids | X239 | silt_100-200cm_Q0p95 | Silt particle proportion (>0.05 mm) in fine soil fraction in g/kg for 100 to 200 cm of the surface – prediction for quantile 0.95 |
| SoilGrids | X240 | soc_0-5cm_mean | Soil organic carbon content in fine soil fraction in dg/kg for 0 to 5 cm of the surface - mean |
| SoilGrids | X241 | soc_0-5cm_Q0p05 | Soil organic carbon content in fine soil fraction in dg/kg for 0 to 5 cm of the surface – prediction for quantile 0.05 |
| SoilGrids | X242 | soc_0-5cm_Q0p5 | Soil organic carbon content in fine soil fraction in dg/kg for 0 to 5 cm of the surface – prediction for quantile 0.5 |
| SoilGrids | X243 | soc_0-5cm_Q0p95 | Soil organic carbon content in fine soil fraction in dg/kg for 0 to 5 cm of the surface – prediction for quantile 0.95 |
| SoilGrids | X244 | soc_5-15cm_mean | Soil organic carbon content in fine soil fraction in dg/kg for 5 to 15 cm of the surface - mean |

| | | | |
|---|---|---|---|
| SoilGrids | X245 | soc_5-15cm_Q0p05 | Soil organic carbon content in fine soil fraction in dg/kg for 5 to 15 cm of the surface – prediction for quantile 0.05 |
| SoilGrids | X246 | soc_5-15cm_Q0p5 | Soil organic carbon content in fine soil fraction in dg/kg for 5 to 15 cm of the surface – prediction for quantile 0.5 |
| SoilGrids | X247 | soc_5-15cm_Q0p95 | Soil organic carbon content in fine soil fraction in dg/kg for 5 to 15 cm of the surface – prediction for quantile 0.95 |
| SoilGrids | X248 | soc_15-30cm_mean | Soil organic carbon content in fine soil fraction in dg/kg for 15 to 30 cm of the surface - mean |
| SoilGrids | X249 | soc_15-30cm_Q0p05 | Soil organic carbon content in fine soil fraction in dg/kg for 15 to 30 cm of the surface – prediction for quantile 0.05 |
| SoilGrids | X250 | soc_15-30cm_Q0p5 | Soil organic carbon content in fine soil fraction in dg/kg for 15 to 30 cm of the surface – prediction for quantile 0.5 |
| SoilGrids | X251 | soc_15-30cm_Q0p95 | Soil organic carbon content in fine soil fraction in dg/kg for 15 to 30 cm of the surface – prediction for quantile 0.95 |
| SoilGrids | X252 | soc_30-60cm_mean | Soil organic carbon content in fine soil fraction in dg/kg for 30 to 60 cm of the surface - mean |
| SoilGrids | X253 | soc_30-60cm_Q0p05 | Soil organic carbon content in fine soil fraction in dg/kg for 30 to 60 cm of the surface – prediction for quantile 0.05 |
| SoilGrids | X254 | soc_30-60cm_Q0p5 | Soil organic carbon content in fine soil fraction in dg/kg for 30 to 60 cm of the surface – prediction for quantile 0.5 |
| SoilGrids | X255 | soc_30-60cm_Q0p95 | Soil organic carbon content in fine soil fraction in dg/kg for 30 to 60 cm of the surface – prediction for quantile 0.95 |
| SoilGrids | X256 | soc_60-100cm_mean | Soil organic carbon content in fine soil fraction in dg/kg for 60 to 100 cm of the surface - mean |
| SoilGrids | X257 | soc_60-100cm_Q0p05 | Soil organic carbon content in fine soil fraction in dg/kg for 60 to 100 cm of the surface – prediction for quantile 0.05 |
| SoilGrids | X258 | soc_60-100cm_Q0p5 | Soil organic carbon content in fine soil fraction in dg/kg for 60 to 100 cm of the surface – prediction for quantile 0.5 |
| SoilGrids | X259 | soc_60-100cm_Q0p95 | Soil organic carbon content in fine soil fraction in dg/kg for 60 to 100 cm of the surface – prediction for quantile 0.95 |
| SoilGrids | X260 | soc_100-200cm_mean | Soil organic carbon content in fine soil fraction in dg/kg for 100 to 200 cm of the surface - mean |
| SoilGrids | X261 | soc_100-200cm_Q0p05 | Soil organic carbon content in fine soil fraction in dg/kg for 100 to 200 cm of the surface – prediction for quantile 0.05 |
| SoilGrids | X262 | soc_100-200cm_Q0p5 | Soil organic carbon content in fine soil fraction in dg/kg for 100 to 200 cm of the surface – prediction for quantile 0.5 |
| SoilGrids | X263 | soc_100-200cm_Q0p95 | Soil organic carbon content in fine soil fraction in dg/kg for 100 to 200 cm of the surface – prediction for quantile 0.95 |
| NasaPower[3] | X264 | T2M 2010a2019ANN | Annual mean temperature at 2 meters above ground for the period 2010 to 2019 |
| NasaPower | X265 | RH2M 2010a2019ANN | Annual mean relative humidity at 2 meters above ground for the period 2010 to 2019 |
| NasaPower | X266 | WS2M 2010a2019ANN | Annual mean wind speed at 2 meters above ground for the period 2010 to 2019 |
| NasaPower | X267 | T2MDEW 2010a2019ANN | Annual mean dew point temperature at 2 meters above ground for the period 2010 to 2019 |
| NasaPower | X268 | T2M MAX 2010a2019ANN | Annual mean maximum temperature at 2 meters above ground for the period 2010 to 2019 |
| NasaPower | X269 | T2M MIN 2010a2019ANN | Annual mean minimum temperature at 2 meters above ground for the period 2010 to 2019 |

| | | | |
|---|---|---|---|
| NasaPower | X270 | PRECTOTCORR 2010a2019ANN | Corrected total precipitation for the period 2010 to 2019. "CORR" indicates that this measurement has been corrected or adjusted in some way |
| NasaPower | X271 | ALLSKY SFC LW DWN 2010a2019ANN | Downward longwave radiation at the surface under all sky conditions for the period 2010 to 2019. Decadal average |
| NasaPower | X272 | ALLSKY SFC SW DWN 2010a2019ANN | Downward shortwave radiation at the surface under all sky conditions for the period 2010 to 2019. Decadal average |
| NasaPower | X273 | alt 2010a2019ANN | Mean altitude of the location where the data were recorded during the period 2010 to 2019 |
| NasaPower | X274 | T2M 2010a2019APR | Mean temperature at 2 meters above ground during April for each year from 2010 to 2019 |
| NasaPower | X275 | RH2M 2010a2019APR | Mean relative humidity at 2 meters above ground during April for the specified years |
| NasaPower | X276 | WS2M 2010a2019APR | Mean wind speed at 2 meters above ground during April over the period 2010 to 2019 |
| NasaPower | X277 | T2MDEW 2010a2019APR | Mean dew point temperature at 2 meters during April for each year in the specified interval. The dew point is an indicator of atmospheric humidity |
| NasaPower | X278 | T2M MAX 2010a2019APR | Mean maximum temperature at 2 meters above ground during April for the years 2010 to 2019 |
| NasaPower | X279 | T2M MIN 2010a2019APR | Mean minimum temperature at 2 meters above ground during April during the same period |
| NasaPower | X280 | PRECTOTCORR 2010a2019APR | Corrected total precipitation recorded during April of each year from 2010 to 2019. Corrections typically adjust various factors to ensure accuracy |
| NasaPower | X281 | ALLSKY SFC LW DWN 2010a2019APR | Downward longwave radiation at the surface under all sky conditions during April of each year in the decade. Measures longwave radiation reaching the Earth's surface |
| NasaPower | X282 | ALLSKY SFC SW DWN 2010a2019APR | Downward shortwave radiation at the surface under all sky conditions during April of each year in the decade. Measures shortwave radiation reaching the Earth's surface |
| NasaPower | X283 | alt 2010a2019APR | Represents the mean altitude of the location(s) where April data were collected throughout those years |
| NasaPower | X284 | T2M 2010a2019AUG | Mean temperature at 2 meters above ground during August for each year from 2010 to 2019 |
| NasaPower | X285 | RH2M 2010a2019AUG | Mean relative humidity at 2 meters above ground during August for the specified years |
| NasaPower | X286 | WS2M 2010a2019AUG | Mean wind speed at 2 meters above ground during August over the period 2010 to 2019 |
| NasaPower | X287 | T2MDEW 2010a2019AUG | Mean dew point temperature at 2 meters during August for each year in the specified interval. The dew point is an indicator of atmospheric humidity |
| NasaPower | X288 | T2M MAX 2010a2019AUG | Mean maximum temperature at 2 meters above ground during August for the years 2010 to 2019 |
| NasaPower | X289 | T2M MIN 2010a2019AUG | Mean minimum temperature at 2 meters above ground during August during the same period |
| NasaPower | X290 | PRECTOTCORR 2010a2019AUG | Corrected total precipitation recorded during August of each year from 2010 to 2019. Corrections typically adjust various factors to ensure accuracy |

| | | | |
|---|---|---|---|
| NasaPower | X291 | ALLSKY SFC LW DWN 2010a2019AUG | Downward longwave radiation at the surface under all sky conditions during August of each year in the decade. Measures longwave radiation reaching the Earth's surface |
| NasaPower | X292 | ALLSKY SFC SW DWN 2010a2019AUG | Downward shortwave radiation at the surface under all sky conditions during August of each year in the decade. Measures shortwave radiation reaching the Earth's surface |
| NasaPower | X293 | alt 2010a2019AUG | Represents the mean altitude of the location(s) where August data were collected throughout those years |
| NasaPower | X294 | T2M 2010a2019DEC | Mean temperature at 2 meters above ground during December for each year from 2010 to 2019 |
| NasaPower | X295 | RH2M 2010a2019DEC | Mean relative humidity at 2 meters above ground during December for the specified years |
| NasaPower | X296 | WS2M 2010a2019DEC | Mean wind speed at 2 meters above ground during December over the period 2010 to 2019 |
| NasaPower | X297 | T2MDEW 2010a2019DEC | Mean dew point temperature at 2 meters during December for each year in the specified interval. The dew point is an indicator of atmospheric humidity |
| NasaPower | X298 | T2M MAX 2010a2019DEC | Mean maximum temperature at 2 meters above ground during December for the years 2010 to 2019 |
| NasaPower | X299 | T2M MIN 2010a2019DEC | Mean minimum temperature at 2 meters above ground during December during the same period |
| NasaPower | X300 | PRECTOTCORR 2010a2019DEC | Corrected total precipitation recorded during December of each year from 2010 to 2019. Corrections typically adjust various factors to ensure accuracy |
| NasaPower | X301 | ALLSKY SFC LW DWN 2010a2019DEC | Downward longwave radiation at the surface under all sky conditions during December of each year in the decade. Measures longwave radiation reaching the Earth's surface. |
| NasaPower | X302 | ALLSKY SFC SW DWN 2010a2019DEC | Downward shortwave radiation at the surface under all sky conditions during December of each year in the decade. Measures shortwave radiation reaching the Earth's surface. |
| NasaPower | X303 | alt 2010a2019DEC | Represents the average altitude of the location(s) where December data were collected throughout those years. |
| NasaPower | X304 | T2M 2010a2019FEB | Mean temperature at 2 meters above ground during February for each year from 2010 to 2019. |
| NasaPower | X305 | RH2M 2010a2019FEB | Mean relative humidity at 2 meters above ground during February for the specified years. |
| NasaPower | X306 | WS2M 2010a2019FEB | Mean wind speed at 2 meters above ground during February over the period from 2010 to 2019. |
| NasaPower | X307 | T2MDEW 2010a2019FEB | Mean dew point temperature at 2 meters during February for each year in the specified interval. The dew point is an indicator of atmospheric humidity. |
| NasaPower | X308 | T2M MAX 2010a2019FEB | Mean maximum temperature at 2 meters above ground during February for the years from 2010 to 2019. |
| NasaPower | X309 | T2M MIN 2010a2019FEB | Mean minimum temperature at 2 meters above ground during February for the same period. |
| NasaPower | X310 | PRECTOTCORR 2010a2019FEB | Total corrected precipitation recorded during February of each year from 2010 to 2019. Corrections typically adjust various factors to ensure accuracy. |

| | | | |
|---|---|---|---|
| NasaPower | X311 | ALLSKY SFC LW DWN 2010a2019FEB | Downward longwave radiation at the surface under all sky conditions during February of each year in the decade. Measures longwave radiation reaching the Earth's surface. |
| NasaPower | X312 | ALLSKY SFC SW DWN 2010a2019FEB | Downward shortwave radiation at the surface under all sky conditions during February of each year in the decade. Measures shortwave radiation reaching the Earth's surface. |
| NasaPower | X313 | alt 2010a2019FEB | Represents the average altitude of the location(s) where February data were collected throughout those years. |
| NasaPower | X314 | T2M 2010a2019JAN | Mean temperature at 2 meters above ground during January for each year from 2010 to 2019. |
| NasaPower | X315 | RH2M 2010a2019JAN | Mean relative humidity at 2 meters above ground during January for the specified years. |
| NasaPower | X316 | WS2M 2010a2019JAN | Mean wind speed at 2 meters above ground during January over the period from 2010 to 2019. |
| NasaPower | X317 | T2MDEW 2010a2019JAN | Mean dew point temperature at 2 meters during January for each year in the specified interval. The dew point is an indicator of atmospheric humidity. |
| NasaPower | X318 | T2M MAX 2010a2019JAN | Mean maximum temperature at 2 meters above ground during January for the years from 2010 to 2019. |
| NasaPower | X319 | T2M MIN 2010a2019JAN | Mean minimum temperature at 2 meters above ground during January for the same period. |
| NasaPower | X320 | PRECTOTCORR 2010a2019JAN | Total corrected precipitation recorded during January of each year from 2010 to 2019. Corrections typically adjust various factors to ensure accuracy. |
| NasaPower | X321 | ALLSKY SFC LW DWN 2010a2019JAN | Downward longwave radiation at the surface under all sky conditions during January of each year in the decade. Measures longwave radiation reaching the Earth's surface. |
| NasaPower | X322 | ALLSKY SFC SW DWN 2010a2019JAN | Downward shortwave radiation at the surface under all sky conditions during January of each year in the decade. Measures shortwave radiation reaching the Earth's surface. |
| NasaPower | X323 | alt 2010a2019JAN | Represents the average altitude of the location(s) where January data were collected throughout those years. |
| NasaPower | X324 | T2M 2010a2019JUL | Mean temperature at 2 meters above ground during July for each year from 2010 to 2019. |
| NasaPower | X325 | RH2M 2010a2019JUL | Mean relative humidity at 2 meters above ground during July for the specified years. |
| NasaPower | X326 | WS2M 2010a2019JUL | Mean wind speed at 2 meters above ground during July over the period from 2010 to 2019. |
| NasaPower | X327 | T2MDEW 2010a2019JUL | Mean dew point temperature at 2 meters during July for each year in the specified interval. The dew point is an indicator of atmospheric humidity. |
| NasaPower | X328 | T2M MAX 2010a2019JUL | Mean maximum temperature at 2 meters above ground during July for the years from 2010 to 2019. |
| NasaPower | X329 | T2M MIN 2010a2019JUL | Mean minimum temperature at 2 meters above ground during July for the same period. |
| NasaPower | X330 | PRECTOTCORR 2010a2019JUL | Total corrected precipitation recorded during July of each year from 2010 to 2019. Corrections typically adjust various factors to ensure accuracy. |

| | | | |
|---|---|---|---|
| NasaPower | X331 | ALLSKY SFC LW DWN 2010a2019JUL | Downward longwave radiation at the surface under all sky conditions during July of each year in the decade. Measures longwave radiation reaching the Earth's surface. |
| NasaPower | X332 | ALLSKY SFC SW DWN 2010a2019JUL | Downward shortwave radiation at the surface under all sky conditions during July of each year in the decade. Measures shortwave radiation reaching the Earth's surface. |
| NasaPower | X333 | alt 2010a2019JUL | Represents the average altitude of the location(s) where July data were collected throughout those years. |
| NasaPower | X334 | T2M 2010a2019JUN | Mean temperature at 2 meters above ground during June for each year from 2010 to 2019. |
| NasaPower | X335 | RH2M 2010a2019JUN | Mean relative humidity at 2 meters above ground during June for the specified years. |
| NasaPower | X336 | WS2M 2010a2019JUN | Mean wind speed at 2 meters above ground during June over the period from 2010 to 2019. |
| NasaPower | X337 | T2MDEW 2010a2019JUN | Mean dew point temperature at 2 meters during June for each year in the specified interval. The dew point is an indicator of atmospheric humidity. |
| NasaPower | X338 | T2M MAX 2010a2019JUN | Mean maximum temperature at 2 meters above ground during June for the years from 2010 to 2019. |
| NasaPower | X339 | T2M MIN 2010a2019JUN | Mean minimum temperature at 2 meters above ground during June for the same period. |
| NasaPower | X340 | PRECTOTCORR 2010a2019JUN | Total corrected precipitation recorded during June of each year from 2010 to 2019. Corrections typically adjust various factors to ensure accuracy. |
| NasaPower | X341 | ALLSKY SFC LW DWN 2010a2019JUN | Downward longwave radiation at the surface under all sky conditions during June of each year in the decade. Measures longwave radiation reaching the Earth's surface. |
| NasaPower | X342 | ALLSKY SFC SW DWN 2010a2019JUN | Downward shortwave radiation at the surface under all sky conditions during June of each year in the decade. Measures shortwave radiation reaching the Earth's surface. |
| NasaPower | X343 | alt 2010a2019JUN | Represents the average altitude of the location(s) where June data were collected throughout those years. |
| NasaPower | X344 | T2M 2010a2019MAR | Mean temperature at 2 meters above ground during March for each year from 2010 to 2019. |
| NasaPower | X345 | RH2M 2010a2019MAR | Mean relative humidity at 2 meters above ground during March for the specified years. |
| NasaPower | X346 | WS2M 2010a2019MAR | Mean wind speed at 2 meters above ground during March over the period from 2010 to 2019. |
| NasaPower | X347 | T2MDEW 2010a2019MAR | Mean dew point temperature at 2 meters during March for each year in the specified interval. The dew point is an indicator of atmospheric humidity. |
| NasaPower | X348 | T2M MAX 2010a2019MAR | Mean maximum temperature at 2 meters above ground during March for the years from 2010 to 2019. |
| NasaPower | X349 | T2M MIN 2010a2019MAR | Mean minimum temperature at 2 meters above ground during March for the same period. |
| NasaPower | X350 | PRECTOTCORR 2010a2019MAR | Total corrected precipitation recorded during March of each year from 2010 to 2019. Corrections typically adjust various factors to ensure accuracy. |

| | | | |
|---|---|---|---|
| NasaPower | X351 | ALLSKY SFC LW DWN 2010a2019MAR | Downward longwave radiation at the surface under all sky conditions during March of each year in the decade. Measures longwave radiation reaching the Earth's surface. |
| NasaPower | X352 | ALLSKY SFC SW DWN 2010a2019MAR | Downward shortwave radiation at the surface under all sky conditions during March of each year in the decade. Measures shortwave radiation reaching the Earth's surface. |
| NasaPower | X353 | alt 2010a2019MAR | Represents the average altitude of the location(s) where March data were collected throughout those years. |
| NasaPower | X354 | T2M 2010a2019MAY | Mean temperature at 2 meters above ground during May for each year from 2010 to 2019. |
| NasaPower | X355 | RH2M 2010a2019MAY | Mean relative humidity at 2 meters above ground during May for the specified years. |
| NasaPower | X356 | WS2M 2010a2019MAY | Mean wind speed at 2 meters above ground during May over the period from 2010 to 2019. |
| NasaPower | X357 | T2MDEW 2010a2019MAY | Mean dew point temperature at 2 meters during May for each year in the specified interval. The dew point is an indicator of atmospheric humidity. |
| NasaPower | X358 | T2M MAX 2010a2019MAY | Mean maximum temperature at 2 meters above ground during May for the years from 2010 to 2019. |
| NasaPower | X359 | T2M MIN 2010a2019MAY | Mean minimum temperature at 2 meters above ground during May for the same period. |
| NasaPower | X360 | PRECTOTCORR 2010a2019MAY | Total corrected precipitation recorded during May of each year from 2010 to 2019. Corrections typically adjust various factors to ensure accuracy. |
| NasaPower | X361 | ALLSKY SFC LW DWN 2010a2019MAY | Downward longwave radiation at the surface under all sky conditions during May of each year in the decade. Measures longwave radiation reaching the Earth's surface. |
| NasaPower | X362 | ALLSKY SFC SW DWN 2010a2019MAY | Downward shortwave radiation at the surface under all sky conditions during May of each year in the decade. Measures shortwave radiation reaching the Earth's surface. |
| NasaPower | X363 | alt 2010a2019MAY | Represents the average altitude of the location(s) where May data were collected throughout those years. |
| NasaPower | X364 | T2M 2010a2019NOV | Mean temperature at 2 meters above ground during November for each year from 2010 to 2019. |
| NasaPower | X365 | RH2M 2010a2019NOV | Mean relative humidity at 2 meters above ground during November for the specified years. |
| NasaPower | X366 | WS2M 2010a2019NOV | Mean wind speed at 2 meters above ground during November over the period from 2010 to 2019. |
| NasaPower | X367 | T2MDEW 2010a2019NOV | Mean dew point temperature at 2 meters during November for each year in the specified interval. The dew point is an indicator of atmospheric humidity. |
| NasaPower | X368 | T2M MAX 2010a2019NOV | Mean maximum temperature at 2 meters above ground during November for the years from 2010 to 2019. |
| NasaPower | X369 | T2M MIN 2010a2019NOV | Mean minimum temperature at 2 meters above ground during November for the same period. |
| NasaPower | X370 | PRECTOTCORR 2010a2019NOV | Total corrected precipitation recorded during November of each year from 2010 to 2019. Corrections typically adjust various factors to ensure accuracy. |

| | | | |
|---|---|---|---|
| NasaPower | X371 | ALLSKY SFC LW DWN 2010a2019NOV | Downward longwave radiation at the surface under all sky conditions during November of each year in the decade. Measures longwave radiation reaching the Earth's surface. |
| NasaPower | X372 | ALLSKY SFC SW DWN 2010a2019NOV | Downward shortwave radiation at the surface under all sky conditions during November of each year in the decade. Measures shortwave radiation reaching the Earth's surface. |
| NasaPower | X373 | alt 2010a2019NOV | Represents the average altitude of the location(s) where November data were collected throughout those years. |
| NasaPower | X374 | T2M 2010a2019OCT | Mean temperature at 2 meters above ground during October for each year from 2010 to 2019. |
| NasaPower | X375 | RH2M 2010a2019OCT | Mean relative humidity at 2 meters above ground during October for the specified years. |
| NasaPower | X376 | WS2M 2010a2019OCT | Mean wind speed at 2 meters above ground during October over the period from 2010 to 2019. |
| NasaPower | X377 | T2MDEW 2010a2019OCT | Mean dew point temperature at 2 meters during October for each year in the specified interval. The dew point is an indicator of atmospheric humidity. |
| NasaPower | X378 | T2M MAX 2010a2019OCT | Mean maximum temperature at 2 meters above ground during October for the years from 2010 to 2019. |
| NasaPower | X379 | T2M MIN 2010a2019OCT | Mean minimum temperature at 2 meters above ground during October for the same period. |
| NasaPower | X380 | PRECTOTCORR 2010a2019OCT | Total corrected precipitation recorded during October of each year from 2010 to 2019. Corrections typically adjust various factors to ensure accuracy. |
| NasaPower | X381 | ALLSKY SFC LW DWN 2010a2019OCT | Downward longwave radiation at the surface under all sky conditions during October of each year in the decade. Measures longwave radiation reaching the Earth's surface. |
| NasaPower | X382 | ALLSKY SFC SW DWN 2010a2019OCT | Downward shortwave radiation at the surface under all sky conditions during October of each year in the decade. Measures shortwave radiation reaching the Earth's surface. |
| NasaPower | X383 | alt 2010a2019OCT | Represents the average altitude of the location(s) where October data were collected throughout those years. |

[1]https://www.worldclim.org/data/index.html;
[2]https://www.isric.org/explore/soilgrids;
[3]https://power.larc.nasa.gov/.

#### == Table S2 ==

**Table S2.** Results from multiple linear mixed models fitted using lmer, including one full model and several reduced versions. The table presents fixed and random effects, Type III ANOVA with Satterthwaite's method for degrees of freedom estimation, variance components, and model comparison metrics (AIC, BIC, LRT).

| **Fixed Effect** | **SumSq** | **MeanSq** | **NumDF** | **DenDF** | **Fvalue** | **p-val (F)** |
|---|---|---|---|---|---|---|
| SITE | 91500132 | 639861 | 143 | 444.05 | 1.5792 | 0.000229*** |

| **Random Effect** | **Variance** | **Std. Dev** | **Variance (in %)** | **Metrics by removing the respective effect** | | | |
|---|---|---|---|---|---|---|---|
| | | | | **AIC** | **BIC** | **LRT ($\chi^2$)** | **p-val ($\chi^2$)** |
| YEAR | 83834.01 | 289.54 | 4.31% | 919676.8 | 921028.5 | 9.494183 | 0.002061244* |
| GEN | 36254.04 | 190.4 | 1.86% | 919771.40 | 921123.00 | 104.03 | 1.99E-24* |
| BLC | 210752.47 | 459.08 | 10.83% | 932013.5 | 933365.20 | 12346.16 | <1.00E-25* |
| GEN:SITE | 26518.9 | 162.85 | 1.36% | 919724.40 | 921076.10 | 57.05 | 4.25E-14* |
| GEN:YEAR | 13260.1 | 115.15 | 0.68% | 919721.90 | 921073.60 | 54.61 | 1.47E-13* |
| SITE:YEAR | 990242.88 | 995.11 | 50.89% | 922161.30 | 923513.00 | 2493.95 | <1.00E-25* |
| GEN:SITE:YEAR | 179882.2 | 424.13 | 9.24% | 922712.50 | 924064.20 | 3045.18 | <1.00E-25* |
| Residual | 405176.88 | 636.54 | 20.82% | – | – | – | – |
| Total | 1945921.48 | – | 100.00% | **919669.30** | **921030.00** | – | – |

| **Model** | **Variance** | **Std. Dev** | **AIC** | **BIC** | **LRT ($\chi^2$)** | **p-val ($\chi^2$)** |
|---|---|---|---|---|---|---|
| MODUNI | 1748527 | 2366.42 | **925987.4** | **927312.3** | 6326.11 | <1.00E-25*** |

For p-values smaller than computational precision, we report <1.00E-25. AIC, BIC, $\chi^2$, and p-values refer to the reduced models (and the MODUNI, the model results used in the article) without the random effect indicated in each row, compared to the full model.

== **Figure S1.** ==

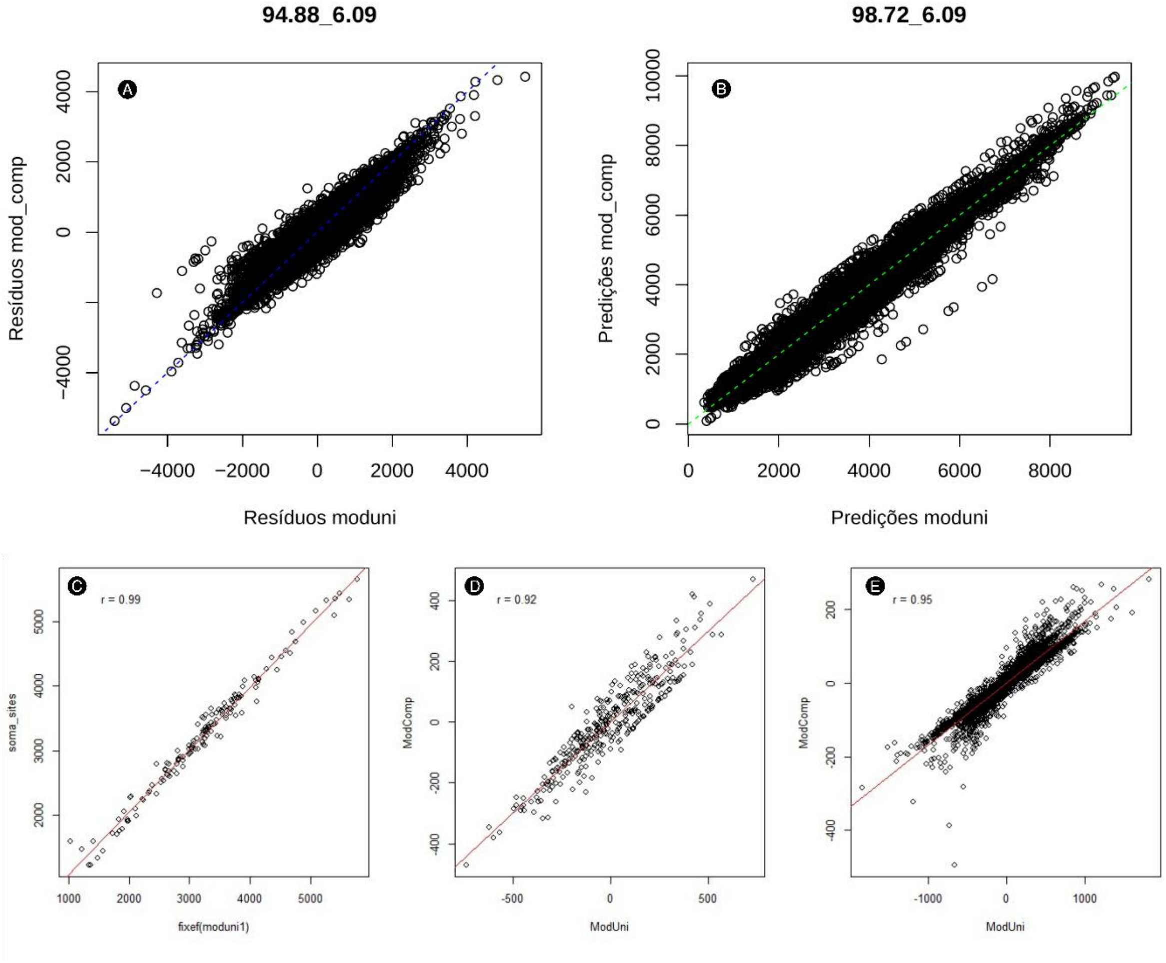


**Figure S1**. **(A)** Residuals from moduni (x-axis) versus residuals from mod_comp (y-axis), with a linear fit in blue. The top label (94.88, 6.09) refers to model metrics. **(B)** Predictions from moduni (x-axis) versus predictions from mod_comp (y-axis), with a linear fit in green. The top label (98.72, 6.09) refers to model metrics. **(C)** Relationship between fixef(moduni) and the fixed effects from the complete model (mod_comp). The red line is a linear fit, and r = 0.99. **(D)** Relationship between gen_moduni intercepts and gen_modcomp intercepts. The red line is a linear fit, and r = 0.92. **(E)** Relationship between intGS_moduni (GEN:SITE) intercepts and intGS_modcomp (GEN:SITE) intercepts. The red line is a linear fit, and r = 0.95.

**== Figure S2. ==**

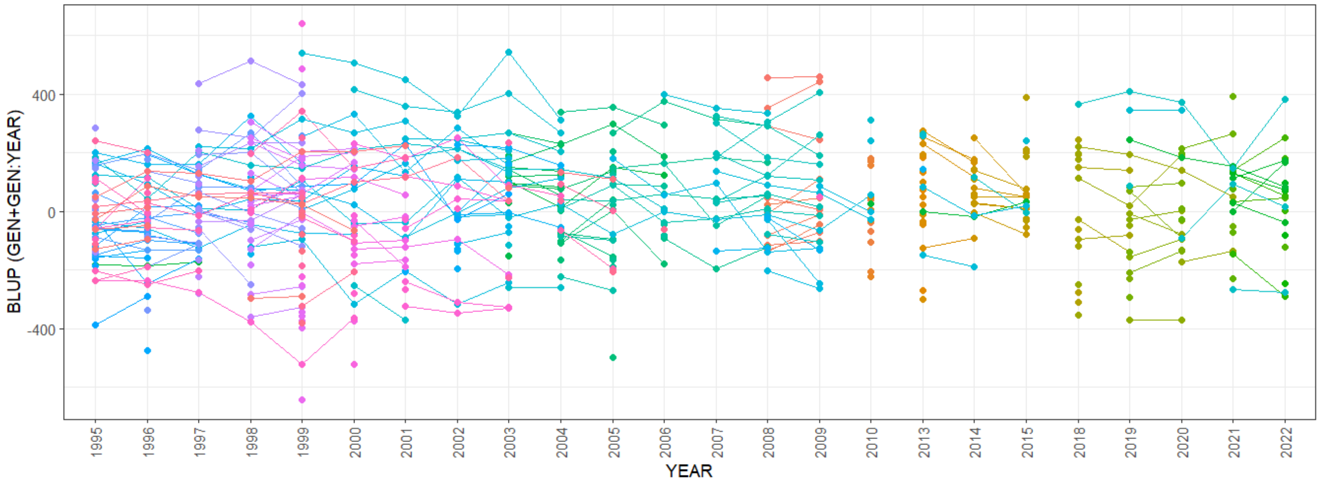


**Figure S2**. BLUP (GEN + GEN:YEAR) values across years from 1995 to 2022, with each color representing a different genotype; points connected by lines indicate repeated measurements for two years in a row.
